# High affinity of Skp to OmpC revealed by single-molecule detection

**DOI:** 10.1101/2020.04.03.022855

**Authors:** Sichen Pan, Chen Yang, Xin Sheng Zhao

## Abstract

Outer membrane proteins (OMPs) are essential to Gram-negative bacteria, and they need molecular chaperones to prevent from aggregation in periplasm during the OMPs biogenesis. Seventeen kilodalton protein (Skp) is the major protein for this purpose. Here we used singlemolecule detection (SMD) to study the stoichiometry modulation of Skp in binding with outer membrane protein C (OmpC) from *Escherichia coli*. To accomplish our task, we developed the tool of portion selectively chosen fluorescence correlation spectroscopy (pscFCS). We found that Skp binds OmpC with high affinity. The half concentration for Skp to form homotrimer Skp_3_ (*C*_1/2_) was measured to be 250 nM. Under the Skp concentrations far below *C*_1/2_ OmpC can recruit Skp monomers to form OmpC·Skp_3_. The affinity of the process is in picomolar range, indicating that the trimerization of Skp in OmpC·Skp_3_ complex is induced by OmpC-Skp interaction even though free Skp_3_ is rarely present. In the concentration range that Skp_3_ is the predominant form, OmpC may directly interact with Skp_3_. Under micro-molar concentrations of Skp, the formation of OmpC·(Skp_3_)_2_ was observed. Our results suggest that the fine-tuned modulation of Skp composition stoichiometry plays an important role in the safe-guarding and quality control mechanism of OMPs in the periplasm.

## Introduction

Outer membrane (OM) in Gram-negative bacteria is crucial for bacterial survival because it separates the periplasm and the external environment and protects cells from toxic molecuels^1^. Meanwhile, most outer membrane proteins (OMPs) adopt a porin-like β-barrel conformation to take nutrients and excrete toxic waste products. As OMPs are synthesized in cytoplasm in unfolded state and have to transport through aqueous periplasm to fold in OM, several periplasmic quality control factors such as survival protein A (SurA), seventeen kilodalton protein (Skp) and serine endoprotease DegP are involved to protect OMPs from mis-folding and aggregation^2–5^. Recent study from our group identifies Skp to be the major protein to prevent OMPs from aggregation^6^. Accumulation of periplasmic OMPs and other survival stress will stimulate the σ^E^ response^7^, which downregulates OMPs expression and upregulates chaperone expression^8^. Crystallographic analysis reveals that Skp exists as a homo-trimer with a “jelly-fish” shape, and the trimeric core is composed of inter-subunit β-sheets and the tentacles of hairpin-shaped α-helices^9,10^. The crystallographic structure makes people believe that Skp_3_ is the basic unit to interact with OMPs. Lyu *et al*. shows that the N-terminus of OMPs enters Skp_3_ firstly through the tentacles and Skp_3_ engulfs entire OMPs via the electrostatic and hydrophobic interactions^11^. The tentacles are remarkably flexible, which modulate the size of cavity to accommodate OMPs with different sizes^12,13^. Meanwhile, only the β-barrel domains of OMPs are captured in the binding cavity^14^ and the encapsulated domains populate a dynamic conformational ensemble^15,16^. The global lifetime of OMPoSkp_3_ complex is hours to ensure OMPs’ searching for low-energy conformations in the cavity^15,17^.

Recently, the dynamic equilibrium between Skp and Skp_3_ has attracted attention. Sandlin *et al*. shows that dynamic equilibrium exists between Skp monomer and Skp_3_ at micro-molar physiological concentration^18^, which is significantly larger than the reported nano-molar affinity of Skp to OMPs^19^. It is therefore speculated that Skp monomer may be involved in the chaperone activity^18^. Moreover, two Skp_3_ are found to bind large OMPs^13^. These studies raise questions on what is the exact composition stoichiometry of Skp in OMPs-Skp complex at low Skp concentration where Skp monomer is the predominant form and under what concentrations these complexes form. Couple obstacles hinder people from getting answers for these questions. Firstly, OMPs are prone to aggregation at ensemble level, which will bring side effect to the experimental phenomenon. Secondly, there are many subpopulations in solution. For example, OMPs are in equilibrium among freely existent (apo-) state and complex formed with chaperones (bound-) state^20^, and Skp has equilibrium between Skp and Skp_3_^18^. These equilibria will blur the information at ensemble level.

To overcome above problems, subpopulation needs to be separately resolved by, for example, electrophoresis, chromatography, or single-molecule detection (SMD)^6,21–23^. Unlike other separation methods which perturb the molecular interaction and the equilibrium of subpopulations, SMD yields a statistical analysis of individual molecule via ultrahigh spatiotemporal resolution. Furthermore, SMD may prepare and investigate OMP samples under monomeric form, so that the interference of OMP aggregation can be completely removed^6,24^. Here we developed an easy-to-implement SMD method named the portion selectively chosen fluorescence correlation spectroscopy (pscFCS), which takes advantages of single-molecule fluorescence resonance energy transfer (smFRET) and fluorescence correlation spectroscopy (FCS) to get diffusion time of specific subpopulation. We used pscFCS as well as fluorescent sodium dodecyl sulfate polyacrylamide gel electrophoresis (SDS-PAGE) to examine the formation and composition stoichiometry of the complexes of Skp and outer membrane protein C (OmpC). We found that under pico-molar concentrations of Skp, OmpC induced the trimerization of Skp to form OmpC·Skp_3_. Under micro-molar concentrations of Skp, additional Skp_3_ can bind OmpC·Skp_3_ to form OmpC·(Skp_3_)_2_. Our results provided new information to help better understanding the role of Skp in the safeguarding and quality control mechanism of OMPs.

## Results

### Development of pscFCS for specific subpopulation

OMPs biogenesis is a complex system, consisting of different subpopulations including OMPs of different states and chaperones of different kinds. To resolve specific subpopulation, we developed pscFCS which utilizes smFRET and FCS^24^. We measured smFRET histogram and chose the fluorescence traces of specific subpopulation (SI methods). Figure 1a shows that under SMD condition the desired fluorescence traces within certain smFRET efficiency portion were selected and the unselected parts of the traces were replaced by random Poisson noise at the experimental level. Figure 1b shows the smFRET histogram before and after the selection on the generated fluorescence traces from Monte-Carlo simulation. In the simulation 6 distinguished molecules diffused freely through focus volume, where 4 molecules (labelled as molecule 1 to 4) had theoretical smFRET efficiency of 0.11 and 2 molecules (labelled as molecule 5 and 6) had theoretical smFRET efficiency of 0.33. The fluorescence traces of smFRET efficiency within the portion of 0.3-0.5, which covered mostly the smFRET efficiency contributed by molecules 5 and 6, were chosen to recalculate the smFRET histogram. The recalculated smFRET histogram only had the smFRET signals from molecule 5 and molecule 6, and removed the contributions of molecule 1 to 4. The synthesized traces of donor channel and acceptor channel were added together to calculate FCS curves, which cancelled the effect of the dynamics between the donor and acceptor and yielded a more accurate diffusion time (Supplementary Figs. S1 and S2). The FCS curve was cut off at 10 μs in the short time edge and was fitted by the 2-dimensional diffusion (2D) model to derive the apparent diffusion time of specific species (SI methods and Supplementary Figs. S3-S8),

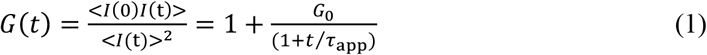

where *I* is the fluorescence intensity, *τ*_app_ is the apparent diffusion time and *G*_0_ is inverse of the number of fluorescent molecules in the focus volume. Obviously, the peak threshold on photon counts in picking up the selected bursts would generate a bias on the apparent diffusion time. Taking Cy3B dye in solution as a real example, the FCS curves of two thresholds are shown in Fig. 1c and the relation between the apparent diffusion time and the peak threshold of three parallel experiments are shown in Fig. 1d. We empirically found that the relation between the apparent diffusion time and the peak threshold can be fitted by

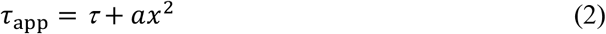

where *x* is the peak threshold, *τ*and *a* are fitting parameters. To correct the bias, an extrapolation procedure was implemented by using equation (2), and *τ* is taken to be the unbiased diffusion time. As a test, the unbiased diffusion time of Cy3B measured by pscFCS was 151±2 μs (n=3, Fig. 1d). While the diffusion time of Cy3B measured by conventional FCS was 162±8 μs (n=5, Supplementary Fig. S9). The relative error was 6.8%, which is sufficient for our goal of identifying the composition stoichiometry of OmpC-Skp complex.

According to the Stokes-Einstein equation, the diffusion coefficient *D* of spherical particles through a liquid with low Reynolds number is

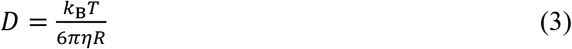

where *η* is the dynamic viscosity and *R* is the hydrodynamic radius. *D* is related to the diffusion time *τ* in FCS by

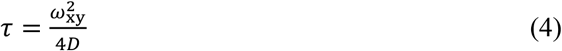

where *ω*_xy_ is the laser beam waist in the *xy* plane. To get molecular weight from *D* by using the Stokes-Einstein equation, we made three reasonable assumptions: 1) Proteins are in spherical shape, 2) the solution is simple so that the Stokes-Einstein equation holds, and 3) all protein species have the same molecular density. Under above assumptions, the following relation holds between two protein species with one of them taken to be a reference,

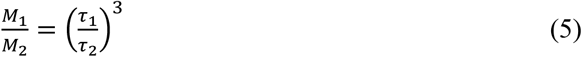

where *M*_1_, *M*_2_, *τ*_1_ and *τ*_2_ are respective molecular weights and diffusion times. The error propagation from the diffusion time to the molecular weight was derived to be

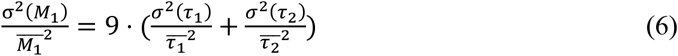

**Figure 1.**
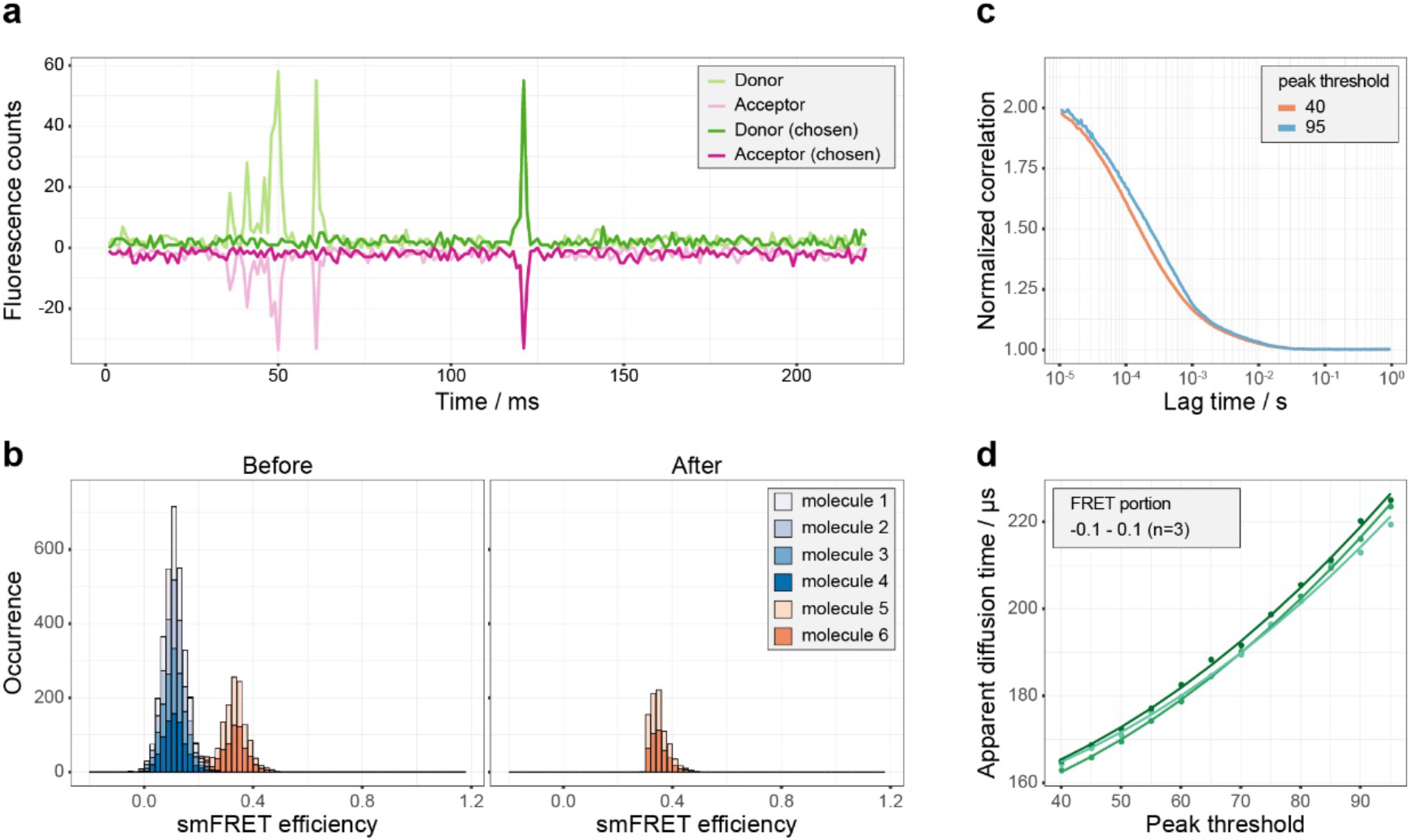
Procedure of yielding unbiased diffusion time from pscFCS for specific subpopulation. (**a**) Fluorescence bursts of interest were chosen and other parts of traces were replaced with Poisson noise. Donor fluorescence bursts before (light green) and after (green) chosen were plotted. Acceptor fluorescence bursts multiplied by −1 before (light magenta) and after (magenta) chosen were plotted. (**b**) the smFRET histogram of the original traces showed two efficiency peaks at the position of 0.11 and 0.33. After selection of smFRET efficiency portion of 0.3-0.5, the recalculated smFRET histogram showed only smFRET efficiency peaks from molecule 5 and 6. (**c**) The peak threshold affected appearance of FCS curve and apparent diffusion time. With larger peak thresholds, the apparent diffusion time of Cy3B was larger. (**d**) Apparent diffusion time showed a non-linear relationship to peak threshold for Cy3B. Dots and lines represent experimental data and fitted curves. Data are shown of three independent experiments.

### Formation of OmpC·Skp_3_ complexes in pM range of Skp concentration

The “jellyfish”-like architecture of Skp_3_ consists of three subunits (Fig. 2a). We first studied the homo-trimerization of Skp by FCS (SI methods and Supplementary Figs. S10 and S11). The dissociation constant *K* was measured to be (4.6±2.7)×10^4^ nM^2^, corresponding to that the half trimerization concentration (*C*_1/2_) of Skp is (2.5±0.7)×10^2^ nM (Supplementary Fig. S12). Our measured *C*_1/2_ is smaller than but on the same order of magnitude with the reported *C*_1/2_ of (4.4±1.3)×10^2^ nM under the most similar temperature and salt concentration^18^, confirming that free Skp is not negligible at a physiological concentration.

To investigate the affinity of Skp to OMPs, we designed intramolecular smFRET experiments, where double mutant OmpC G8C-D335C was labelled by fluorescent dyes AF555 and AF647 and incubated with various concentrations of Skp. Because smFRET efficiency reflects the conformational changes of OmpC between apo-state and bound-state^6,25^, different apparent smFRET efficiencies (*E*_app_) correspond to different species. Figure 2b shows smFRET histogram of OmpC G8C-D335C in presence of certain Skp concentrations. When Skp was absent, only the *E*_app_=0.78 peak was observed. At Skp concentration of 0.23 nM, an smFRET efficiency peak at *E*_app_=0.13 was observed besides the 0.78 peak. When Skp was 1.8 nM, only the 0.13 peak remained (more results in Supplementary Fig. S13). The 0.78 peak was assigned to apo-OmpC and the 0.13 peak to bound-OmpC as previously observed^6^. The peak at *E*_app_=0 (zero peak) was due to missing or inactivated acceptors and was disregarded.

To determine the composition stoichiometry of OmpC-Skp complexes, we used pscFCS to compare the diffusion time of apo- and bound-OmpC in the same smFRET histogram (Fig. 2c). The stoichiometric ratio *r* = Skp: OmpC was derived to be

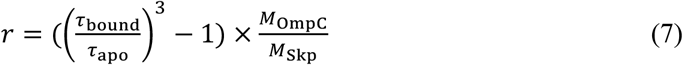

and was determined to be 2.8±0.4 with the apo-OmpC molecule as the inner reference, indicating that OmpC was already bound by Skp_3_ even though the concentration of Skp in solution was extremely lower than *C*_1/2_.

To accurately quantify the affinity of Skp to OmpC, we conducted colocalization measurement using total internal reflection fluorescence (TIRF) microscope where OmpC G8C-AF555 was immobilized on surface and incubated with freely diffusing Skp D128C-AF647. The concentration of Skp generating half occupation of OmpC (*K*_D_) for the reaction

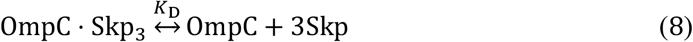

was measured to be (5.5±0.4)×10^2^ pM with a Hill coefficient *n*_Hill_ = 1.6 ± 0.2 (Fig. 3a, SI method). Our measured *K*_D_ is much smaller than previously reported values^19^, which may be due to the high sensitivity of the SMD method. The statistical analysis on the 640 nm-excited fluorescence counts showed that in the OmpC-Skp complex the composition stoichiometry of Skp is larger than 1 (Fig. 3b and Supplementary Fig. S14), consistent with the pscFCS results that there are 3 Skp molecules in the OmpC-Skp complex.

**Figure 2.**
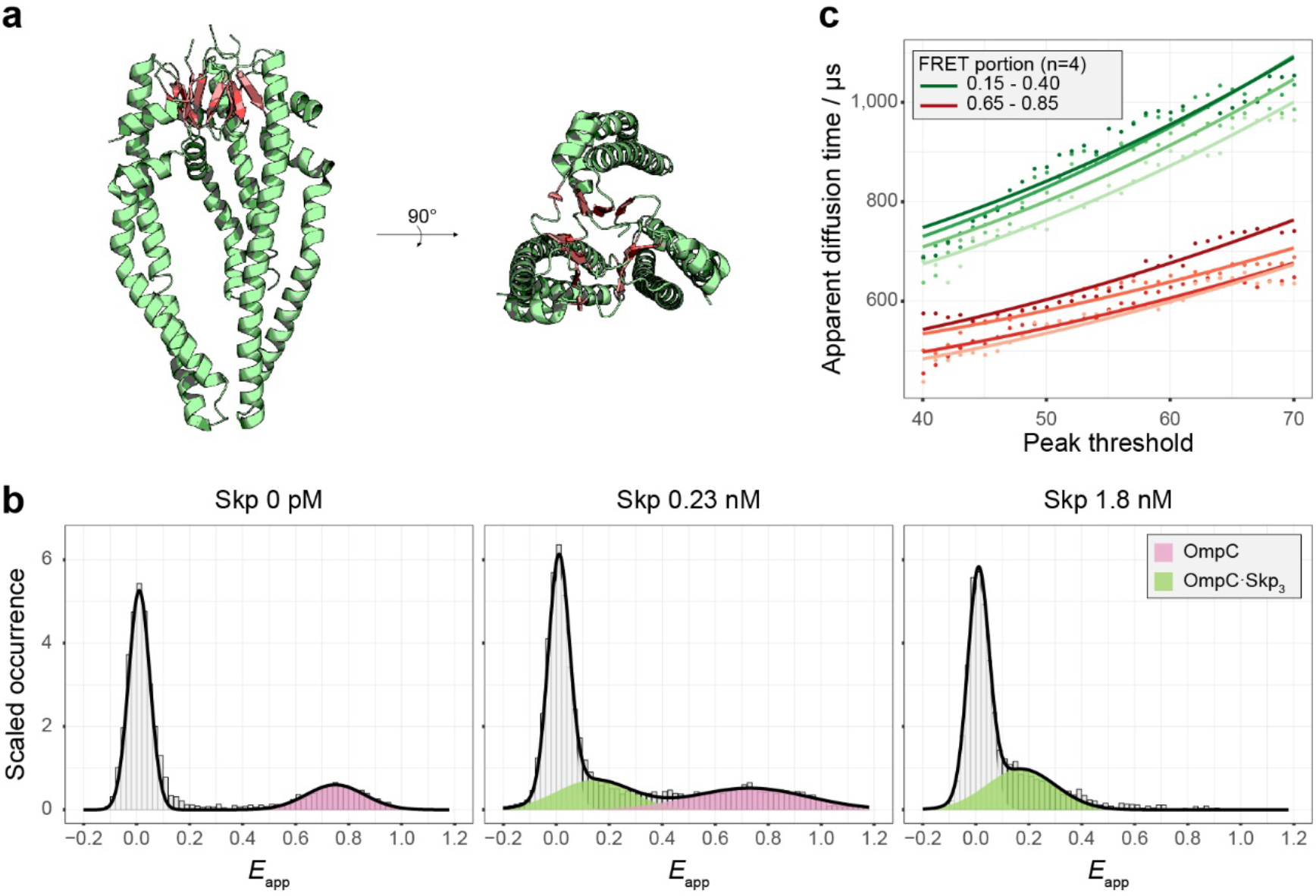
Observation of OmpC·Skp_3_ complexes in pM Skp concentration. (**a**) the crystal structure of Skp_3_ (PDB ID: 1SG2^9^) is shown in side-view and top-view. The β-sheets of each monomer are coloured in salmon and composed of the core domain of Skp_3_. (**b**) The smFRET histogram showed the conformation changes of OmpC G8C-D335C incubated with various concentrations of Skp. These smFRET histograms exhibited two smFRET peaks positioned at 0.13 (green) and 0.73 (pink), which represented bound- and apo-OmpC respectively. Zero-efficiency peak was resulted from missing or inactivated acceptors. All histograms were normalized and fitted by Gaussian distribution. (**c**) pscFCS at different peak thresholds of the bound- (green, FRET portion 0.15 - 0.40) and apo- (red, FRET portion 0.65 - 0.85) OmpC were treated. The positive bias on the apparent diffusion time caused by the peak threshold was corrected by equation (2). The diffusion times of 549±27 μs for the bound- and 422±26 μs for the apo-OmpC were derived. Dots and lines represent experimental data and fitted curves. Data are shown of four independent experiments.

**Figure 3.**
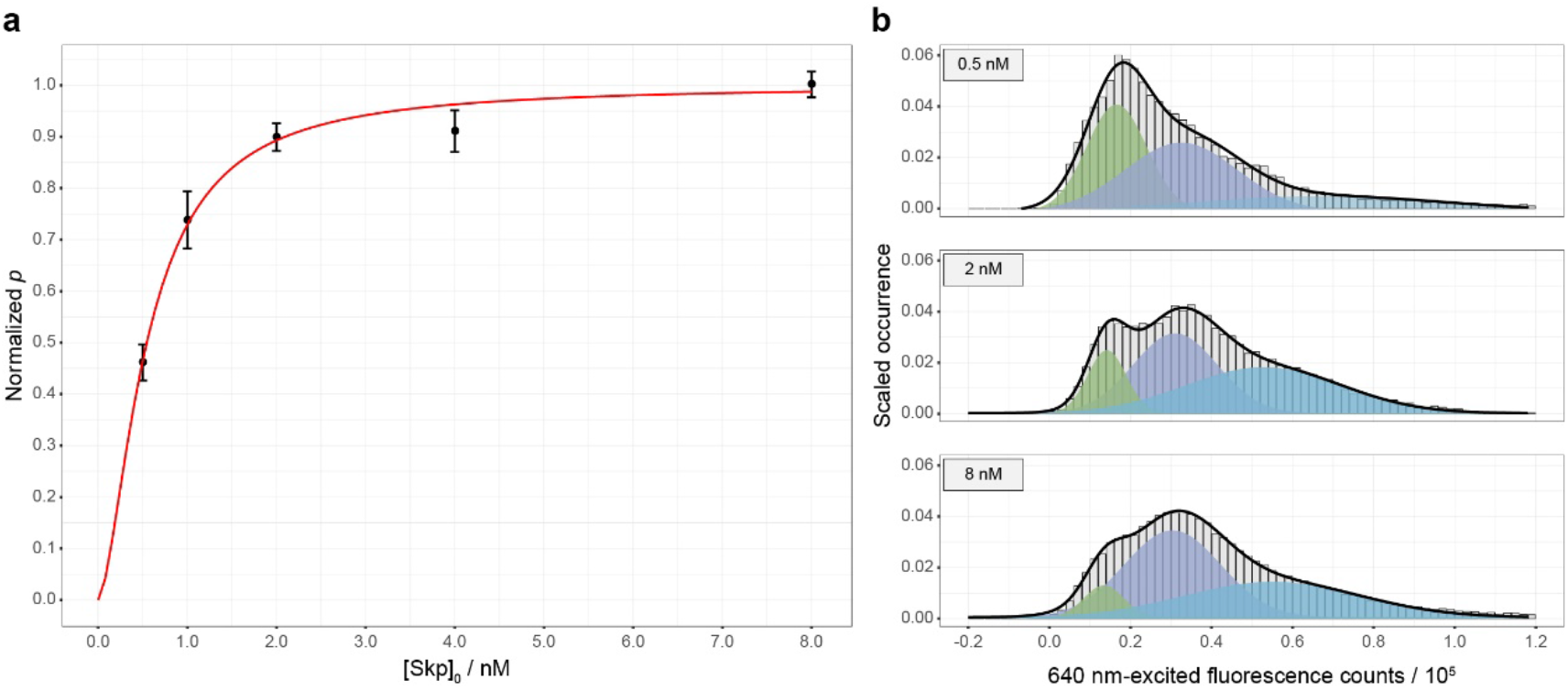
Quantification of affinity of Skp to OmpC. **(a)** The normalized event counts of colocalized donor-acceptor pairs on TIRF experiments (normalized *p*) as a function of total added Skp concentration ([Skp]o) was fitted by Supplementary equation (19) to get the dissociation constant. Black dots and red line represent experimental data and fitted curve. Data are shown as mean±s.d. of six imaging area. **(b)** 640 nm-excited fluorescence counts histogram exhibited three peaks respectively positioned at 0.15×10^5^ (green), 0.31×10^5^ (indigo) and 0.56×10^5^ (azure), which demonstrated that the composition stoichiometry of Skp in the OmpC-Skp complex is larger than 1. All histograms were normalized and fitted by Gaussian distribution.

### Formation of OmpC·(Skp_3_)_2_ at Skp concentrations in μM magnitude

As is shown in Fig. 4a and Supplementary Fig. S13, when intramolecular labelled OmpC G8C-D335C was incubated with micro-molar of Skp, the *E*_app_=0.3 peak (orange) appeared in the smFRET histogram besides previously seen 0.1 peak (green). The phenomenon was checked among different OmpC double mutants (Supplementary Figs. S15-S17). Previous study points out that Skp_3_ is multivalent^13^, so an intuitive hypothesis was that the conformational change of OmpC under micro-molar of Skp was due to the composition stoichiometry change of the OmpC-Skp complexes. To verify the hypothesis, OmpC incubated with micro-molar of Skp was amine-crosslinked and separated in SDS-PAGE (Fig. 4b). When OmpC G8C-AF647 and Skp D128C-Cy3B was mixed, six co-localized gel bands of Cy3B and AF647 from 60 kD to 180 kD were observed. As shown in Fig. 4b, the control experiments showed that when only Skp was present Skp_3_ could be crosslinked intramolecularly but not intermolecularly, and when only OmpC was present in 8 M urea buffer and crosslinked, no intermolecular crosslinked product was observed. The six colocalized gel bands were assigned from OmpC·Skp to OmpC·Skp6 respectively according to their molecular weight (40.2 kD for His-tagged OmpC G8C-AF647 and 18.8 kD for His-tagged Skp D128C-Cy3B). The determined molecular weights of the assigned bands were slightly larger than the theoretical molecular weights due to the association of DSS molecules (Supplementary Table S1). Because of the false negative and false positive possibilities, crosslinked products do not necessarily represent the real species in solution, but it suggests the possible existence of OmpC·Skp6 in the solution. Then, the intramolecular labelled OmpC G8C-D335C and Skp was crosslinked (Supplementary Fig. S18) and the gel bands of OmpC·Skp_3_ and OmpC·Skp6 were excised and retrieved back into buffer to measure their smFRET efficiencies (Fig. 4c). The histogram showed that the retrieved OmpC·Skp_3_ had an *E*_app_ peak at 0.1 and the retrieved OmpC·Skp6 had an *E*app peak at 0.3, consistent with the observed peaks under *in situ* condition (Fig. 4a). The result confirmed the hypothesis that the conformational change of OmpC under micro-molar of Skp was due to the composition stoichiometry change of the OmpC-Skp complexes. By fitting the variation of the peak area in the smFRET histogram under *in situ* condition against the Skp concentration, the dissociation constant *K*_D_ for the reaction

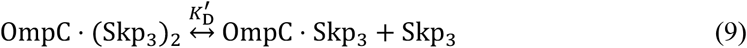

was measured to be 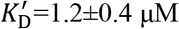 (Fig. 4d, SI method).

Interestingly, the pscFCS determined that the hydrodynamic radius of OmpC·(Skp_3_)_2_ was comparable to the hydrodynamic radius of OmpCSkp_3_ (Supplementary Fig. S19 and Supplementary Table S2), indicating that OmpC·(Skp_3_)_2_ adopts an “inter-locked” configuration (see discussion section for the detail).

**Figure 4.**
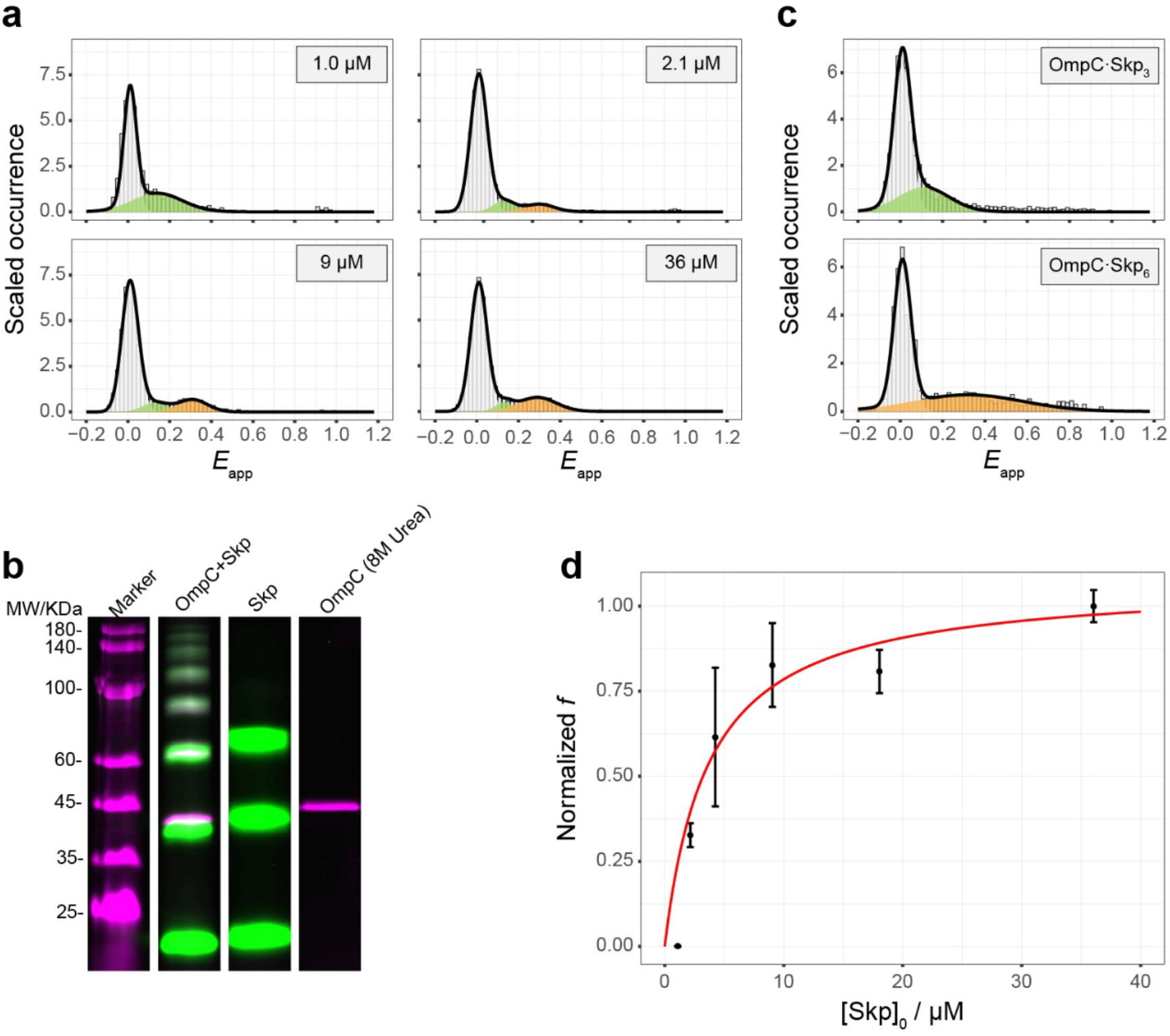
Observation of OmpC(Skp_3_)_2_ complex in μM Skp concentrations. (**a**) As Skp concentration increased, smFRET histogram of OmpC G8C-D335C showed *E*_app_=0.1 peak (green) decreased and *E*_app_=0.3 peak (orange) increased. All histograms were normalized and fitted by Gaussian distribution. (**b**) OmpC-AF647 (magenta) and Skp-Cy3B (green) were incubated, crosslinked and separated by fluorescent SDS-PAGE, which showed six colocalized gel bands of OmpC-Skp⊓ complexes (n=1 to 6). In the negative control of Skp or OmpC alone, no intermolecular crosslinked product was observed. (**c**) intramolecular labelled OmpC G8C-D335C was incubated with Skp and the gel bands of OmpC·Skp_3_ and OmpC·Skp6 were excised and retrieved back into buffer to measure their smFRET efficiencies. smFRET peaks of OmpC·Skp_3_ (green) and OmpC·Skp6 (orange) were identical to that under *in situ* condition respectively. All histograms were normalized and fitted by Gaussian distribution. **(d)** The normalized fraction of [OmpC·(Skp_3_)_2_] over [OmpCSkp_3_]+[OmpC·(Skp_3_)_2_] (normalized *f*) as a function of [Skp]_0_ was fitted by Supplementary equation (21). Black dots and red line represent experimental data and fitted curve. Data are shown as mean±s.d. of three experiments.

## Discussion

We made thorough study on the equilibrium constants of Skp homo-trimerization and the complex formation between OmpC and Skp. Skp D128C was used for fluorescent dye labelling. Although the residue D128 is at trimeric core of Skp_3_ structure, our previous study^11^ shows that the Skp mutation and dye labelling does not perturb its chaperone activity, and labelled Skp D128C had a minimal self-quenching effect. Since OmpC is unfolded, the mutation and dye labelling do not vary its property^6^. The *C*_1/2_ of Skp homo-trimerization was 250 nM, which is comparable with previous measurement^18^, suggesting that Skp monomer cannot be neglected in its biological function as chaperones. When OMP polypeptide was synthesized by ribosome in the cytoplasm, the polypeptide would be secreted to periplasm through SecYEG/SecA translocon in unfolded state and safeguarded by chaperones including Skp_3_^1,26–29^. Many articles showed that OMPoSkp_3_ has a dissociation constant of nano-molar range^19,30,31^. Most their experiments, however, were performed at ensemble level of OMPs, where chaperones had to compete with OMPs self-aggregation. In our smFRET experiments, monodisperse OmpC was prepared which removed the side effect of aggregation^6^, and it was a better mimic of the situation in living cells. With SMD, we were able to resolve individual subpopulations, and we showed high affinity (*K*_D_ = (5.5 ± 0.4) × 10^2^pM) and positive cooperativity (*n*_Hill_ = 1.6 ± 0.2) of Skp to form OmpC·Skp_3_ (Fig. 2 and Fig. 3). Because the immobilized OmpC was limited in phase space compared to freely diffusing OmpC, the affinity in TIRF experiment could be slightly weaker than that in smFRET experiment in solution^32^, but it still resulted in high affinity of Skp towards its client, providing strong evidence to support the proposed unique chaperone function of Skp to dissolve aggregated OmpC^6^. The high affinity of Skp to OmpC is perhaps a result of multiple, non-specific and transient interactions^11,15^. Additionally, Skp has a broad substrate spectrum^33^. We speculate that the high affinity of Skp to OmpC is likely to be prevalent among its other substrates.

The reaction rate of Skp binding OmpC is nearly diffusion limited^30^. The association rate constant could be estimated by

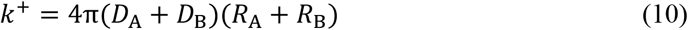

where *D* is the diffusion coefficient, and *R* is the radius of molecules. The radius of Skp_3_ in solution is 3.3 nm^34^, and the radius of Skp is 2.3 nm derived from our FCS data. According to equations (3) and (10), the association rate constant of Skp to OmpC is 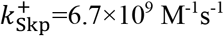, and Skp_3_ to OmpC is 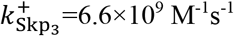. When the equilibrium between Skp and Skp_3_ is considered, the apparent reaction rate constants are defined as 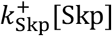 and 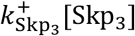. The apparent reaction rate constant of Skp is several orders of magnitude larger than that of Skp_3_ in low Skp concentrations. With the increase of the Skp concentration, the apparent reaction rate constant of Skp_3_ becomes progressively predominant (Supplementary Fig. S20). Therefore, in the Skp low concentration range, the induced Skp trimerization by OmpC should be dominant, while the direct reaction between Skp_3_ and OmpC will become the major route when the concentration of Skp_3_ exceeds certain amount. At even higher Skp concentrations, OmpC·(Skp_3_)_2_ can form.

Periplasm is lack of ATP as energy source. The OMPs folding free energy of −18 to −32 kcalomol^−1^ acts as an energy sink in the OMPs biogenesis^31^. Our results revealed that the free energy of Skp_3_ binding to OmpC is −20 kcal-mol^−1^, which supports the ideas that Skp_3_ is a superior holdase^17^ and de-aggregation agency^6^ to OMPs. Since the binding energy is so large, the release of the substrates from Skp may need the help of LPS^35,36^, DegP^2,37^ or BAM system^38,39^, ensuring the direction of OMPs transportation towards the final destination.

We observed the formation of OmpC·(Skp_3_)_2_, but the hydrodynamic radius of OmpC·(Skp_3_)_2_ was similar to that of OmpC·Skp_3_ according our pscFCS results (Supplementary Table S2). This experimental result strongly suggests that the orientation of the two Skp_3_ in OmpC·(Skp_3_)_2_ is via the “inter-locked” pattern^13^. Our data showed that the intramolecular smFRET efficiency of OmpC is higher in OmpC·(Skp_3_)_2_ than that in OmpC·Skp_3_, indicating that OmpC is compressed tighter in OmpC·(Skp_3_)_2_ than in OmpC·Skp_3_, that also causes the reduction of the complex radius and is consistent with the “inter-locked” model. SANS study reveals that the gyration radius of apo-state Skp_3_ in solution is larger than Skp_3_ bound to OmpA or OmpW^34^. OmpA and OmpW are both small OMPs with 8 ß-strands. It is reasonable that the complex is larger when Skp_3_ binds to larger OMPs such as OmpC (16 ß-strands). When apo-state Skp_3_ in solution was used as reference, we found that indeed the hydrodynamic radius of OmpC·Skp_3_ is larger than that of Skp_3_.

The physiological concentration of Skp is around 2.1-3.9 μM at stationary phase growth in LB^6,18^, which is near the dissociation constant 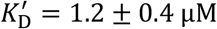 for second Skp_3_. The σ^E^ response will downregulate OMPs expression and upregulate expression level of chaperones and proteases^40^. Although the whole OMPs biogenesis landscape will be much complicated *in vivo*, we speculate that the upregulated Skp will increase the population of OMP·(Skp_3_)_2_, which may enhance the protection of OMPs from aggregation to help the cell survival. Figure 5 presents a diagram how the composition stoichiometry in the OmpC-Skp complex is proceeded as the Skp concentration varies. The fine-tuned modulation of different complex forms may play a distinct role in the OMPs biogenesis under different situations. Our results provide new insight regarding the mechanism of Skp’s chaperone function in OMPs biogenesis.

**Figure 5.**
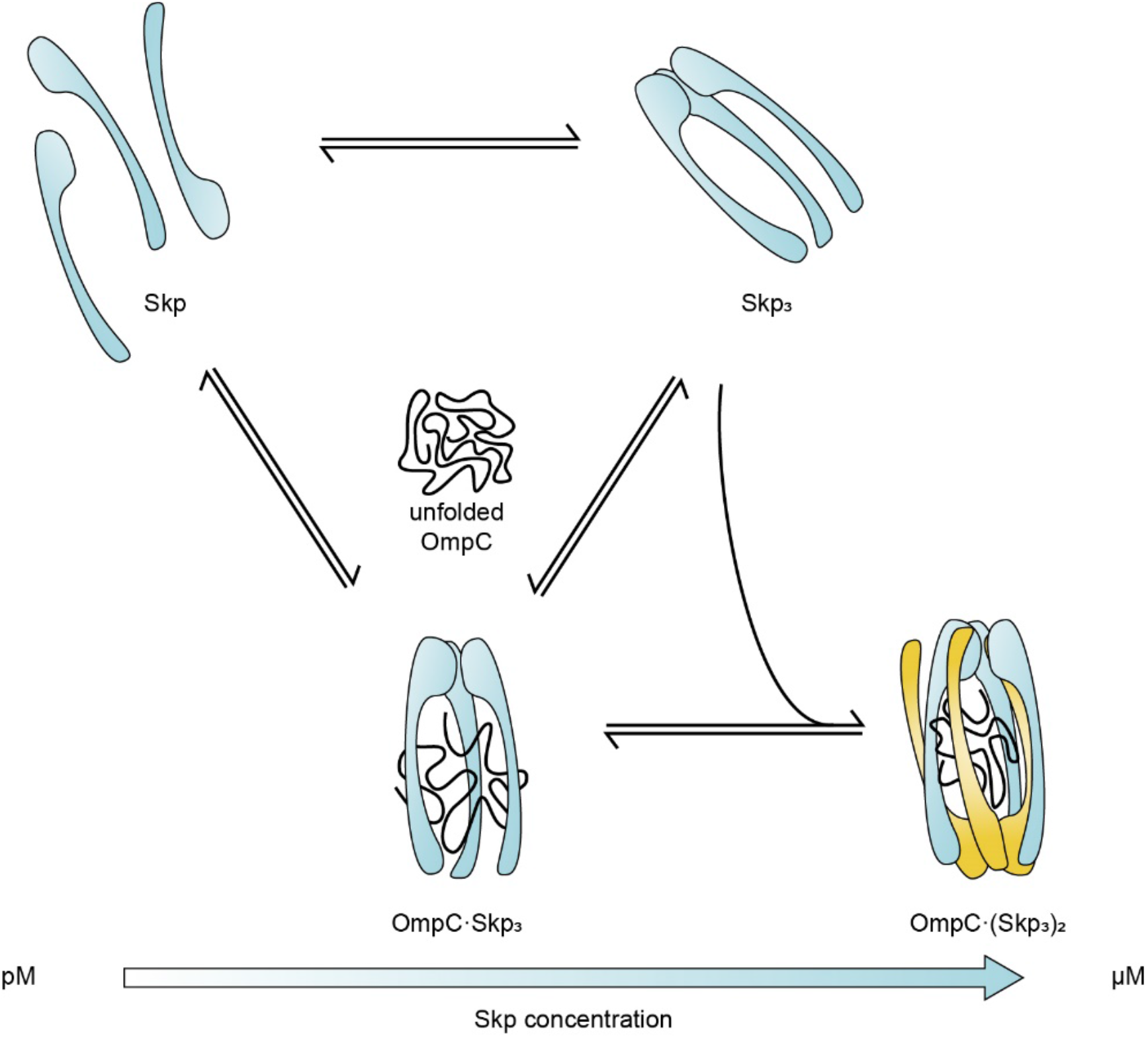
Schematic view of Skp binding OmpC. OmpC collapses in aqueous solution without chaperones. When pico-molar of Skp is present, Skp binds OmpC to form OmpC·Skp_3_ complex. In sub micro-molar concentrations, Skp_3_ may directly react with OmpC to form OmpC·Skp_3_. When micro-molar of Skp is present, OmpC·(Skp_3_)_2_ will appear with an “interlocked” binding pattern.

## Methods

### Protein expression, purification and mutagenesis

The pET28a vectors carrying relevant Skp, including N-terminal His-tag, were transformed into *E.coli*. BL21(DE3) pLysS cells (TransGen Biotech). Cells were grown in LB medium containing 50 μg/mL kanamycin at 37 °C with shaking (220 r.p.m.) until the culture reached an OD_600_ of ~0.6 (after ~3 h). The culture was induced with 0.5 mM IPTG, expressed for 4 h and harvested by centrifugation. The cell pellet was resuspended in 80 mL buffer A (50 mM PB, 500 mM NaCl and 10 mM imidazole, pH=8.0) and lysed by ultrasonication. The supernatant was collected by centrifugation at 20,000 g for 30 min, filtered by 0.22 μm syringe filter and loaded on Ni-NTA (5 mL) column (GE Healthcare) equilibrated with buffer A. The column was washed extensively by buffer A and eluted with a gradient (25%, 100%) of buffer B (50 mM PB, 500 mM NaCl and 250 mM imidazole, pH=8.0). Protein fractions were dialyzed against buffer C (50 mM PB and 100 mM NaCl, pH=7.0) by desalting column (GE Healthcare), and concentrated by centrifugal filter units (Merck Amicon). Aliquots were snap-frozen in liquid nitrogen and stored in −80 °C. For the purification of OmpC, the plasmids including N-terminal His-tag were transformed into *E.coli*. BL21(DE3) cells (TransGen Biotech) and 8 M urea was included in buffers A-C to prevent aggregation. For immobilization, an AviTag (GLNDIFEAQKIEWHE) was introduced to the C-terminus of OmpC G8C. Skp concentration was determined by BCA protein assay kit (PIERCE), according to manufacturer’s instructions. Skp monomer was taken in the calculation of the Skp concentrations. OmpC concentration was determined by using molar extinction coefficient of 65,210 M^−1^cm^−1^ at 280 nm.

Fast Mutagenesis System (TransGen Biotech) with plasmids pET28a-*skp* and pET28a-*ompC* as templates was used to construct the site-directed cysteine mutants of Skp and OmpC. For intramolecular smFRET assays, OmpC G8C-L139C, OmpC G8C-I232C and OmpC G8C-D335C were generated. For TIRF colocalization assays, OmpC G8C with AviTag on C-terminus and Skp D128C were generated. Skp D128C was also used in FCS for homo-trimerization assay. The protein expression and purification were the same as described above.

### Fluorescent dye labelling

Labelling of site-directed cysteine mutants of Skp and OmpC was performed as follows. 10fold molar excess of tris-2-carboxyethyl-phosphinie (TCEP) was used in proteins of 100 μM at 27 °C for 30 min to reduce cysteine residues. 5-fold molar excess of maleimide fluorescent dyes Cy3B (GE Healthcare), AF555 (Life Technology) or AF647 (Life Technology) was added and the solution was kept in the dark at 27 °C for 3 h. Excess dyes was removed by HiTrap desalting column (GE Healthcare) in buffer C. 4 M urea was included in the labelling and eluting buffer for Skp to disassemble trimers and expose cysteine residues. 8 M urea was included in the labelling and eluting buffer for OmpC to prevent aggregation. The absorbance of Cy3B (130,000 M^−1^cm^−1^), AF555 (158,000 M^−1^cm^−1^) and AF647 (239,000 M^−1^cm^−1^) was used to determine the fluorescent dye concentration. For all samples, the extent of labelling was near 90%.

### Biotinylation of dye-labelled OmpC

The bioitinylation of OmpC was performed by enzyme BirA, which can specifically recognize AviTag’s lysine side chain and attach biotin. Biotinylation reaction was performed in accordance with the instructions provided by the BirA enzyme kit (GeneCopoeia™). Reaction was conducted including 40 μM dye-labelled OmpC with AviTag, 50 μM D-Biotin, 10 mM ATP, 10 mM MgOAc and 45 U/μL BirA in buffer C containing 4 M urea at 30 °C for 1 hour. After reaction aliquots were snap-frozen in liquid nitrogen and stored in −80 °C.

### Monte-Carlo simulation for Brownian motion

Monte-Carlo simulation was performed as described previously^41,42^ and simplified by putting 6 diffusing molecules into the volume. Each fluorescence trace was tracked and calculated to obtain smFRET histogram where contribution from each molecule was known. Then all fluorescence traces were added to simulate the signals, which was used to select fluorescence bursts of certain subpopulation in pscFCS, and the generated Boolean array was used in each fluorescence trace to test the contribution of 6 molecules by checking selected fluorescence bursts and recalculating smFRET histogram.

### FCS experiments for homo-trimerization of Skp

Homo-trimerization titration of Skp was carried out with a home-built inverted fluorescence confocal microscope based on a TE2000-U microscope (Nikon) equipped with a 532 nm solid-state laser (MLL-III-532-20mW, LD&TEC) as previously described^25,43^. The laser was filtered to 100 μW and focused inside the sample solution through an oil immersion objective (NA 1.45, 60×, Nikon). The fluorescence was separated from the excitation light by a dichromic mirror (zt532 rdc, Chroma). After being focused through a 50 μm pinhole, the fluorescence was separated by a polarizing beam splitter (PBS) (Daheng, China) into reflected s-polarized and transmitted p-polarized beams. The beams were focused on two avalanche photon diodes (APD) (SPCM-AQRH-14, Perkin-Elmer Optoelectronics) respectively for FCS measurement. For each sample, 30 nM of Skp D128C-Cy3B was added in various concentrations of Skp denatured by 4 M urea. The solution was diluted 100-fold in buffer C and incubated in the dark at 24 °C for 30 min to assemble. The final concentration of Skp D128C-Cy3B was 300 pM. 40 μL of solution was sealed between a Secure-Seal™ hybridization chamber gasket (Life Technology) and a cover glass. Experiments were conducted at 23 °C. Surface adsorption of proteins was prevented by 0.02% (v/v) tween 20 (Surfact-Amps 20, Life Technology) in buffer. Then the apparent diffusion time *τ* of sample was obtained by fitting the measured FCS curve using a formula considering 2-dimentional diffusion plus one exponential relaxation (2D1R):

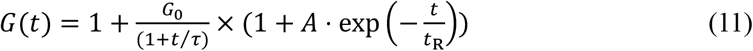

where *τ* is the diffusion time, *t*_R_ is the relaxation time, Gθ is the inverse of number of fluorescent molecules in the focus volume and *A* is the amplitude of the relaxation. The *τ* was used to calculate effective Skp composition stoichiometry *n* according to the Stokes-Einstein equation, where diffusion time and molecular weight of Cy3B were used as reference:

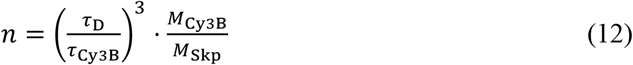

where *M*_Cy3B_ and *M*_skp_ are molecular weights of Cy3B and Skp, and *τ*_Cy3B_ is the diffusion time of Cy3B. The *n* as a function of [Skp]o was fitted by Supplementary equation (11) - (15) to obtain the dissociation constant for Skp homo-trimerization.

### smFRET measurement

smFRET experiments were carried out by using the same confocal microscope. After the pinhole, the fluorescence was divided by a dichroic mirror T635 lpxr (Chroma) into donor and acceptor channels with APD. BrightLine 593/40 nm (Semrock) was put before APD in donor channel and HQ 685/40 nm (Chroma) was put before APD in acceptor channel. 40 μL of solution was sealed between a Secure-Seal™ hybridization chamber gasket (Life Technology) and a cover glass. For intramolecular smFRET, 8 M urea denatured dual-labelled OmpC was diluted into buffer solution having desired Skp concentration, and the final concentration of OmpC was 50 pM. Samples were incubated in the dark for 15 min at room temperature and experiments were conducted at 23 °C. Surface adsorption of proteins was prevented by 0.02% (v/v) tween 20 (Surfact-Amps 20, Life Technology). Fluorescence of every sample was collected for 30 min in 1 ms bintime. Fluorescence data was processed by Python scripts to yield smFRET efficiency by

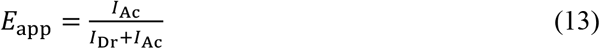

where *E*_app_ is the apparent smFRET efficiency, *I*_Dr_ and *I*_Ac_ are photon counts of every identified fluorescence burst. Statistics of smFRET efficiency yielded smFRET histogram and the histogram was fitted by Gaussian distributions.

### pscFCS measurement for stoichiometry

pscFCS experiments were performed as the same of smFRET experiments except that the bintime was taken to be 0.96 μs. The data of every sample was processed by Python scripts to obtain unbiased diffusion time, which was used to derive stoichiometry by

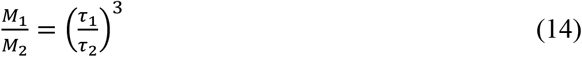

where *M*_1_, *M*_2_, *τ*_1_ and *τ*_2_ are the molecular weights and corrected diffusion times of subpopulations. More details about pscFCS is in supplementary information.

### TIRF colocalization measurement for affinity

The PEG-passivated slides were prepared as previously described^44^. OmpC G8C-AF555 with AviTag on C-terminus was immobilized on coverslip surface via biotin-streptavidin interaction. The cell was incubated by 0.05 mg/mL streptavidin and washed by buffer C. Then 8 M urea denatured biotinylated OmpC was diluted 50-fold in buffer C to final concentration of 50 pM and added to the cell with streptavidin. Then the cell was washed again by buffer C. Finally varying Skp D128C-AF647 from 500 pM to 8 nM in buffer C was added to the cell. Surface adsorption of proteins was prevented by 0.02% (v/v) tween 20 (Surfact-Amps 20, Life Technology) in buffer and an oxygen scavenging system was included in buffer during detection^44^. Colocalization measurement was performed on a home-built TIRF microscope using alternating laser excitation (ALEX) between 532 nm and 640 nm at 23 °C^44^. Emitted fluorescence from the molecules on surface was separated by filters and collected by a dualview EMCCD with frame frequency of 10 Hz. Colocalized fluorescence spots of donor and acceptor excited by 532 nm laser were counted as described before^44^. Traces of monomeric OmpC bound by Skp were selected to yield 640 nm-excited fluorescence counts histogram and apparent smFRET efficiency histogram. Apparent smFRET efficiency was calculated by equation (13).

### Cross-linked fluorescent SDS-PAGE and retrieved single-molecule FRET

For the cross-linked fluorescent SDS-PAGE analysis, samples were prepared by 50-fold dilution of OmpC G8C-AF647 into 0.6 μM of Skp with 0.3 μM of Skp D128C-Cy3B in buffer C, and the final concentration of OmpC G8C-AF647 was 45 nM. Then the sample was incubated at room temperature in the dark for 15 min before adding cross linker DSS (Thermo Fisher) with a final concentration of 20 μM. The sample was incubated for another 30 min, separated by SDS-PAGE and fluorescent imaged by Typhoon FLA 9500 (GE Healthcare). For the retrieved smFRET experiments, samples were incubated and cross-linked as the same procedures as described above. Then 1 M glycine was added to a final concentration of 10 mM to quench the cross-linking reaction. The sample was concentrated by centrifugal filter units and separated by SDS-PAGE. The corresponding gel bands were excised and lysed in 200 μL buffer C at 4 °C, 220 rpm shaking overnight. Then the sample was centrifugated and the supernatant was purified by centrifugal filter units with 10-fold volume of buffer C. Finally, the sample was diluted in buffer C to single-molecule concentration to carry out the smFRET experiments.

### Data Availability

The datasets and code in the study are available from the corresponding author on reasonable request.

## Acknowledgements

We thank Ms. S. Peng and Prof. C. Chen of Tsinghua University for their kind assistance with the TIRF experiment. The project was supported by NSFC (21521003 and 21233002) and NKBRSF (2012CB917304).

## Authors Contributions

S.C.P. and X.S.Z. conceived this study. S.C.P. performed simulation, crosslinked SDS-PAGE experiments and single-molecule fluorescence experiments. C.Y. performed biotinylated sample preparation. S.C.P. and X.S.Z analysed experimental results and wrote the paper. All authors reviewed the manuscript.

