## supporting information for "High affinity of Skp to OmpC revealed by single-molecule detection"

### Supplementary Notes

#### Supplementary note 1: Data processing in pscFCS

First, the high-resolution (0.96 μs-bintime) fluorescence traces of sample were recorded. The traces were binned to 1.008 ms bintime to generate normal single-molecule fluorescence resonance energy transfer (smFRET) histogram. Next, we summed the original high-resolution fluorescence traces of donor and acceptor channels together and selected the portions that contain the whole fluorescence bursts belonging to specific subpopulation in the smFRET histogram. The rest parts of the original high-resolution fluorescence traces were replaced by Poisson noise. Then we used the synthesized high-resolution traces to calculate and fit the FCS curves, which yielded the apparent diffusion time of the specific subpopulation.

Fluorescence fluctuation due to the singlet-triplet transition or other relaxations may have impact on extracting the diffusion time. To evaluate the effect due to fluorescence resonance energy transfer (FRET) process, we simulated 20 sets of two component Brownian motion with an accompanying FRET process of different relaxation times. The auto-correlations of acceptor trace only, donor trace only and trace of acceptor and donor summed together were compared. The FCS curves were fitted by either 2-dimentional diffusion with 1 relaxation term (2D1R) model,

$$G(t) = 1 + \frac{G_0}{(1+\frac{t}{\tau_{\mathrm{app}}})} \times \left(1 + A \cdot \exp\left(-\frac{t}{t_{\mathrm{R}}}\right)\right) \qquad (1)$$

or 3-dimentional diffusion with 1 relaxation term (3D1R) model,

$$G(t) = 1 + \frac{G_0}{\left(1+\frac{t}{\tau_{\mathrm{app}}}\right)\left(1+\frac{t}{\tau_{\mathrm{app}}\cdot\omega^2}\right)^{\frac{1}{2}}} \times \left(1 + A \cdot \exp\left(-\frac{t}{t_{\mathrm{R}}}\right)\right) \qquad (2)$$

where $\tau_{\mathrm{app}}$ is the apparent diffusion time, $t_{\mathrm{R}}$ is the relaxation time, $G_0$ is the inverse of numbers of fluorescent molecules in the focus volume, $A$ is the amplitude for the relaxation and $\omega$ is the the ratio of beam waist in the $z$ direction to beam waist in the $xy$ plane.

From the simulation we found that in the auto-correlation of acceptor trace or donor trace, the apparent diffusion time decreased when the relaxation exists, and the apparent diffusion time depended on specific relaxation dynamics of molecules. However, the auto-correlation of the summed traces was not influenced by the existence of the FRET relaxation (Supplementary Fig. S1). We also found that the 2D1R fitting yielded systematically smaller diffusion time than the theoretical values, whereas the diffusion time of the 3D1R fitting matched the theoretical values well. However, since we used reference to scale the diffusion time, the difference between the ratios on the diffusion time extracted from the 2D1R or 3D1R model was not significant (Supplementary Fig. S2).

Fluorescence count threshold (named as peak threshold) was taken to identify the fluorescence bursts. With higher peak threshold, higher fluorescence bursts were chosen. FCS of higher bursts would yield larger apparent diffusion time because the molecules were deeper in the Gaussian-shaped focus volume,

$$I = I_0 \mathrm{e}^{-2\left(\frac{x^2}{\omega_{\mathrm{xy}}^2}+\frac{y^2}{\omega_{\mathrm{xy}}^2}+\frac{z^2}{\omega_{\mathrm{z}}^2}\right)} \qquad (3)$$

where $I_0$ is the laser intensity at the center, $\omega_{xy}$ is the beam waist in the $xy$ plane and $\omega_z$ is the beam waist in the $z$ direction. An empirical polynomial equation was used to get unbiased diffusion time,

$$\tau_{app} = \tau + ax^2 \tag{4}$$

where $x$ is the peak threshold, $\tau$ is the unbiased diffusion time and $a$ is a fitting parameter. Our simulations demonstrated that the first-order derivative of $\tau_{app}$ at the origin was zero. Therefore, we did not include the linear term in the equation.

In the example of 50 pM of freely diffusing Cy3B, the FRET portion of -0.1 to 0.1 (Supplementary Fig. S3) was chosen to calculate FCS series at different peak threshold.

We also used Cy3B to examine which model better reduced the effect of the singlet-triplet transition. The transition usually occurs on μs timescale, so we compared the results of 10 μs-started FCS curves fitted by a 2-dimensional diffusion (2D) model,

$$G(t) = 1 + \frac{G_0}{(1+t/\tau_{app})} \tag{5}$$

with the results of 1 μs-started FCS curves fitted by 2D or 2D1R models. For the 10 μs-started FCS curves, the singlet-triplet transition was mostly dropped out, so the 2D model was suitable to fit (Supplementary Fig. S4) and yielded apparent diffusion times of 151±2 μs as shown in Fig. 1d in the main text. For the 1 μs-started FCS curves, the singlet-triplet transition was obvious, and the 2D model could not fit the curves well (Supplementary Fig. S5). The extrapolated unbiased diffusion time (Supplementary Fig. S6) was 143±2 μs. For the

2D1R, although the 1 μs-started FCS curves could be well fitted (Supplementary Fig. S7), the apparent diffusion time was irregular due to large fitting errors (Supplementary Fig. S8). The diffusion time of Cy3B measured by conventional FCS was 162±8 μs (Supplementary Fig. S9). Therefore, we chose 2D model to fit 10 μs-started FCS curves of the summed traces in our pscFCS.

**Supplementary note 2: Derivation of dissociation constant for Skp homo-trimerization**

The Skp homo-trimerization is described by

$$\mathrm{Skp_3} \overset{K}{\leftrightarrow} \mathrm{3Skp} \tag{6}$$

$$K = \frac{[\mathrm{Skp}]^3}{[\mathrm{Skp_3}]} \tag{7}$$

where $K$ is the dissociation constant. When dye-labelled Skp is used for measurement while the concentration of unlabelled Skp far exceeds that of dye-labelled Skp, the trimerization with $\mathrm{Skp}^*$ as a component becomes

$$\mathrm{Skp_3^*} \overset{K^*}{\leftrightarrow} \mathrm{Skp^*} + \mathrm{2Skp} \tag{8}$$

$$K^* = \frac{[\mathrm{Skp^*}][\mathrm{Skp}]^2}{[\mathrm{Skp_3^*}]} \tag{9}$$

where $\mathrm{Skp}^*$ is the dye-labelled Skp, $\mathrm{Skp_3^*}$ is Skp trimer which contains one dye-labelled Skp monomer, and $K^*$ is the respective dissociation constant.

FCS experiments measured the effective diffusion time $\tau$ of the species containing dye-labelled Skp. Then it can be converted to the molecular weight and then to the effective composition stoichiometry of the detected Skp species ($n$), which is the average of the numbers of Skp monomer in Skp$^*$ ($n_m$) and $\mathrm{Skp}_3^*$ ($n_t$) over the concentrations of $\mathrm{Skp}^*$ and $\mathrm{Skp}_3^*$:

$$n = n_t \cdot \frac{[\mathrm{Skp}_3^*]}{[\mathrm{Skp}_3^*]+[\mathrm{Skp}^*]} + n_m \cdot \frac{[\mathrm{Skp}^*]}{[\mathrm{Skp}_3^*]+[\mathrm{Skp}^*]} \qquad (10)$$

where $n_t$=3 and $n_m$=1 by definition. Substituting equation (9) into equation (10) we have

$$n = n_t \cdot \left(\frac{[\mathrm{Skp}]^2}{K^*+[\mathrm{Skp}]^2}\right) + n_m \cdot \left(\frac{K^*}{K^*+[\mathrm{Skp}]^2}\right) \qquad (11)$$

Since $[\mathrm{Skp}^*] \ll [\mathrm{Skp}]$, the consumption of Skp by reaction (8) can be neglected and the mass balance can be written as

$$[\mathrm{Skp}]_0 = [\mathrm{Skp}] + 3[\mathrm{Skp}_3] \qquad (12)$$

where $[\mathrm{Skp}]_0$ is the total added concentration of unlabelled Skp. Combining equations (7) and (12), we derive that

$$[\mathrm{Skp}]^3 + \frac{K}{3} \cdot [\mathrm{Skp}] - \frac{K}{3} \cdot [\mathrm{Skp}]_0 = 0 \qquad (13)$$

which has a single real root as

$$[\mathrm{Skp}] = \left(\frac{K}{6} \cdot [\mathrm{Skp}]_0 + (\Delta)^{\frac{1}{2}}\right)^{\frac{1}{3}} + \left(\frac{K}{6} \cdot [\mathrm{Skp}]_0 - (\Delta)^{\frac{1}{2}}\right)^{\frac{1}{3}} \qquad (14)$$

where $\Delta$ is

$$\Delta = \left(\frac{\frac{K}{3}\cdot[\mathrm{Skp}]_0}{2}\right)^2 + \left(\frac{K}{9}\right)^3 \tag{15}$$

The $n$ as a function of $[\mathrm{Skp}]_0$ can be fitted by substituting equation (14) into equation (11) to obtain both the dissociation constants $K^*$ and $K$. The fraction of Skp as $\mathrm{Skp}_3$ over $[\mathrm{Skp}]_0$ ($y$) is

$$y = \frac{3[\mathrm{Skp}_3]}{[\mathrm{Skp}]_0} \tag{16}$$

Substituting equation (12) and equation (16) into equation (13), the relation between $[\mathrm{Skp}]_0$ and $y$ is derived to be

$$[\mathrm{Skp}]_0 = \left(\frac{K}{3}\cdot\frac{y}{(1-y)^3}\right)^{\frac{1}{2}} \tag{17}$$

In the experiment to measure the homo-trimerization equilibrium constant between Skp and $\mathrm{Skp}_3$, we labelled fluorescent dye Cy3B to mutant Skp D128C. Skp D128C-Cy3B at concentration of 0.3 nM was mixed with Skp ranged from 0 to $10^3$ nM in buffer C (50 mM PB, 100 mM NaCl, pH 7.0). FCS was measured on home-built confocal microscope and data was fitted by the 2D1R model (Supplementary Fig. S10). The titration curve of $n$ as a function of $[\mathrm{Skp}]_0$ was plotted and fitted by using above equations (Supplementary Fig. S11). This titration curve showed that $\mathrm{Skp}^*$ kept monomer until around $[\mathrm{Skp}]_0$=37 nM. The dissociation constant $K$ was determined to be $(4.6\pm2.7)\times10^4$ $\mathrm{nM}^2$, indicating that half-trimer concentration ($C_{1/2}$) was of $(2.5\pm0.7)\times10^2$ nM (Supplementary Fig. S12).

**Supplementary note 3: Derivation of equilibrium constants for the formation of OmpC-Skp complexes**

For the equilibrium

$$\mathrm{OmpC}\cdot\mathrm{Skp_3} \overset{K_\mathrm{D}}{\leftrightarrow} \mathrm{OmpC} + 3\mathrm{Skp} \tag{18}$$

where $K_\mathrm{D}$ is the concentration of Skp generating half occupation of OmpC or the apparent dissociation constant . The normalized event counts of colocalized donor-acceptor pairs on TIRF experiments (*p*) is fitted by

$$p = \frac{[\mathrm{Skp}]_0{}^{n_\mathrm{Hill}}}{K_\mathrm{D}^{n_\mathrm{Hill}} + [\mathrm{Skp}]_0{}^{n_\mathrm{Hill}}} \tag{19}$$

where $[\mathrm{Skp}]_0$ is the total concentration of Skp in solution and $n_\mathrm{Hill}$ is Hill coefficient representing the cooperativity of reaction. Normalized *p* against $[\mathrm{Skp}]_0$ was plotted and fitted to obtain $K_\mathrm{D}$ and $n_\mathrm{Hill}$ (Fig. 3a in the main text).

For the equilibrium

$$\mathrm{OmpC}\cdot(\mathrm{Skp_3})_2 \overset{K'_\mathrm{D}}{\leftrightarrow} \mathrm{OmpC}\cdot\mathrm{Skp_3} + \mathrm{Skp_3} \tag{20}$$

where $K'_\mathrm{D} = \frac{[\mathrm{OmpC}\cdot\mathrm{Skp_3}][\mathrm{Skp_3}]}{[\mathrm{OmpC}\cdot(\mathrm{Skp_3})_2]}$ is the respective dissociation constant. The fraction of $[\mathrm{OmpC}\cdot(\mathrm{Skp_3})_2]$ over $[\mathrm{OmpC}\cdot\mathrm{Skp_3}]+[\mathrm{OmpC}\cdot(\mathrm{Skp_3})_2]$ (*f*) represented by the respective smFRET peak areas is derived to be

$$f = \frac{[\mathrm{Skp}]_0}{3K'_\mathrm{D} + [\mathrm{Skp}]_0} \tag{21}$$

Fittings of smFRET peak areas in Supplementary Fig. S13 was carried out to obtain *f*. Then, normalized *f* as a function of $[\text{Skp}]_0$ was plotted and fitted to obtain $K'_{\text{D}}$ (Fig. 4d in the main text).

**Supplementary Figures**

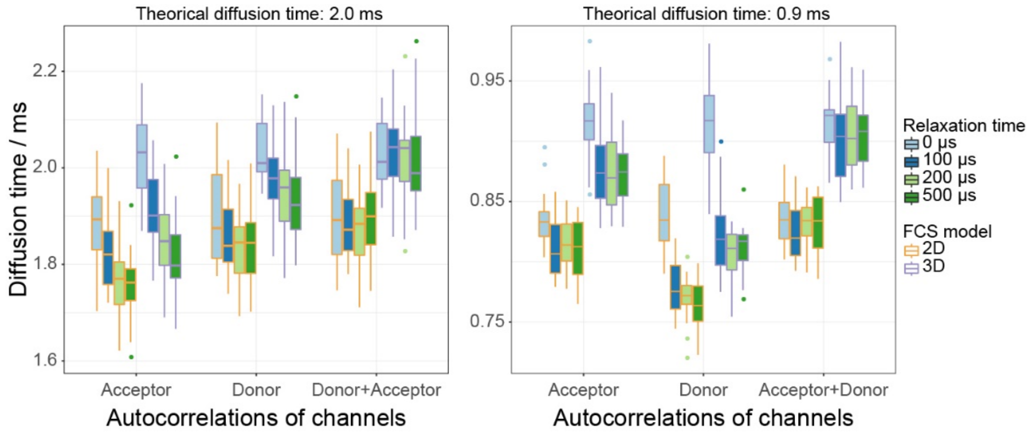


**Supplementary Figure S1** Influence of relaxation time and fashion of donor/acceptor correlations on the extracted diffusion time in the simulated pscFCS.

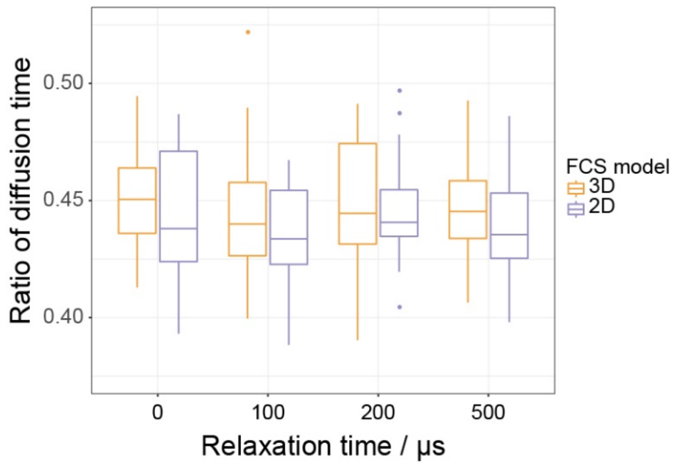


**Supplementary Figure S2** Comparison of the 2D model to the 3D model on the ratio of extracted diffusion times of two species with the theoretical diffusion times of 0.9 ms and 2.0

ms respectively.

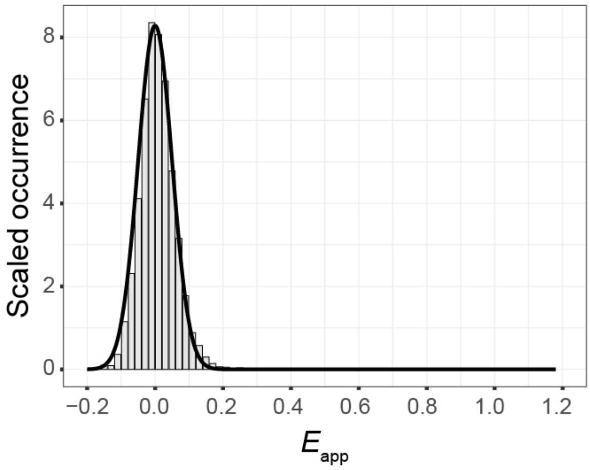


**Supplementary Figure S3** smFRET histogram of Cy3B. The histogram only has zero-efficiency peak due to absence of acceptor dyes. FRET portion of -0.1 to 0.1 was used to calculate pscFCS curves.

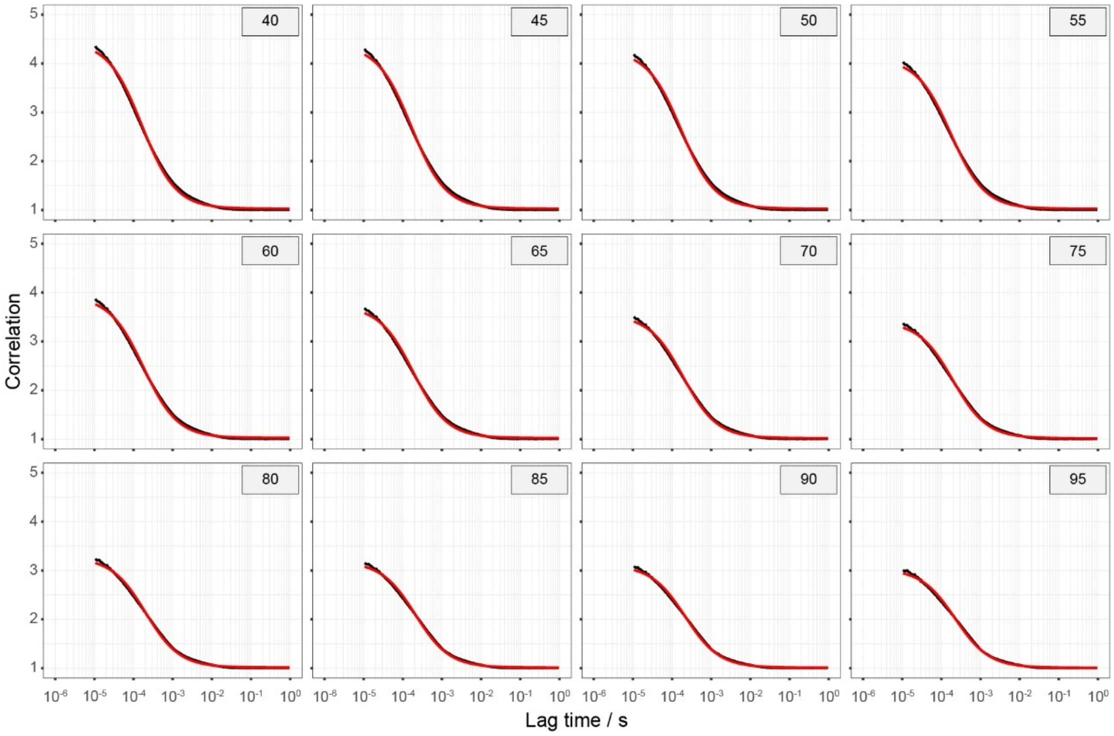


**Supplementary Figure S4** FCS curves starting from 10 μs and respective 2D fit of Cy3B

(FRET portion -0.1 - 0.1) at different peak thresholds. Black line and red line represent experimental data and fitted curves. The extracted apparent diffusion time series were plotted and fitted in Fig. 1d in the main text.

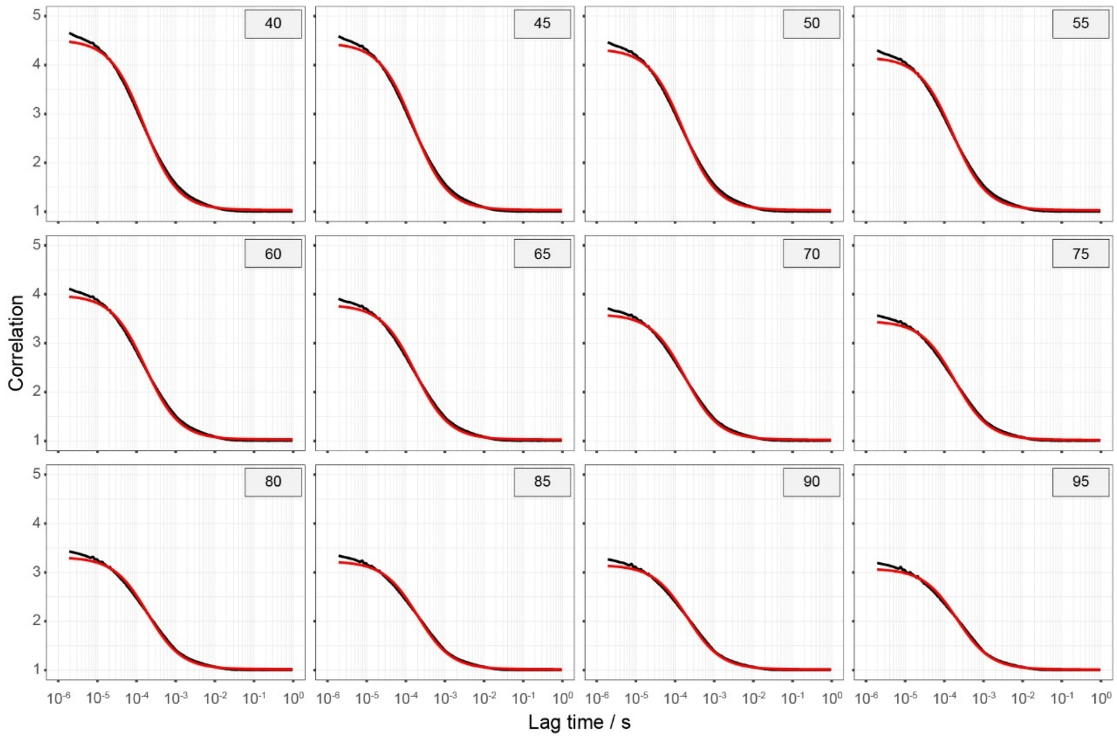


**Supplementary Figure S5** FCS curves starting from 1 μs and respective 2D fit of Cy3B (FRET portion -0.1 - 0.1) at different peak thresholds. Black line and red line represent experimental data and fitted curves.

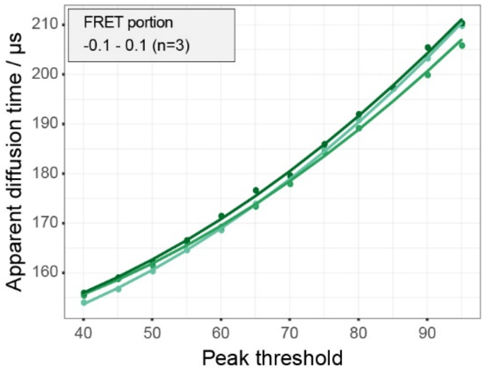


**Supplementary Figure S6** Plot of the apparent diffusion time extracted from the 2D fit of the 1 µs-started FCS curves against the peak threshold. The unbiased diffusion time was 143±2 µs. Dots and lines represent experimental data and fitted curves. Data are shown of three independent experiments.

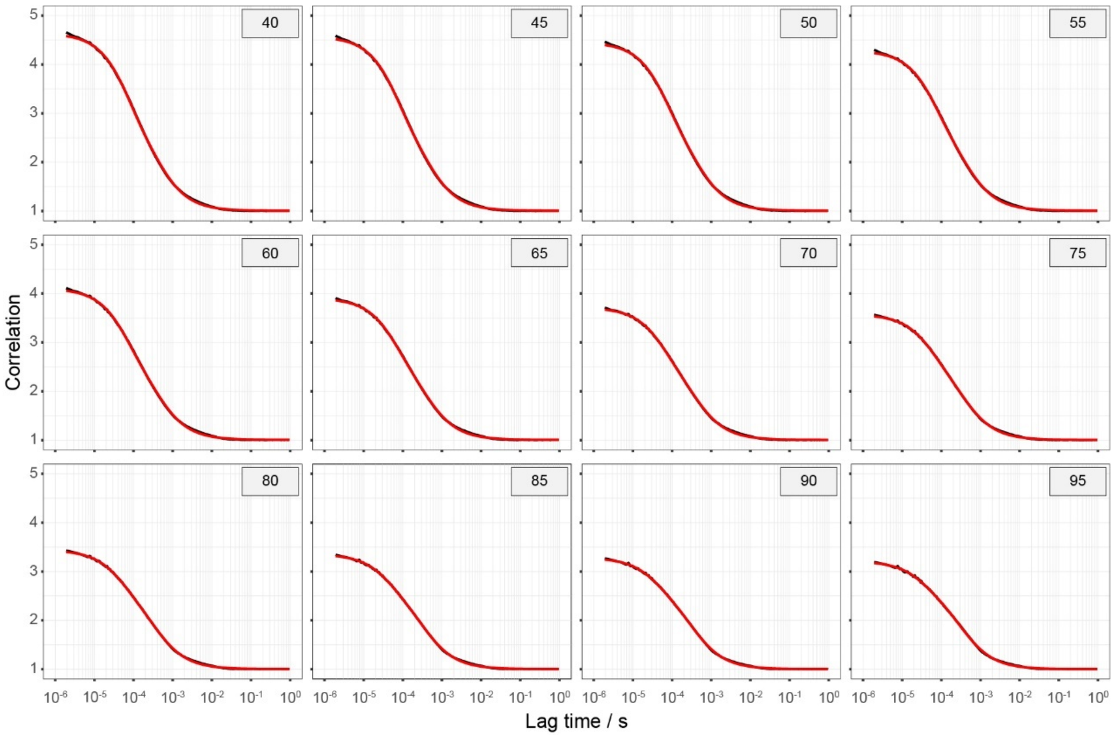


**Supplementary Figure S7** FCS curves starting from 1 µs and respective 2D1R fit of Cy3B (FRET portion -0.1 - 0.1) at different peak thresholds. Black line and red line represent experimental data and fitted curves.

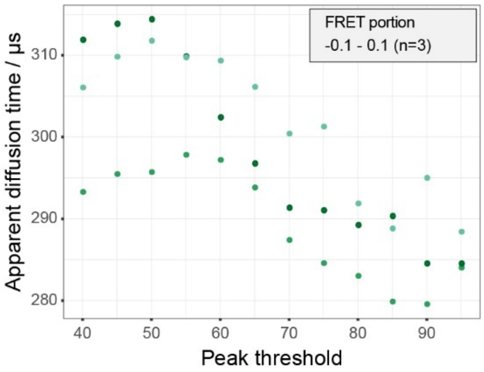


**Supplementary Figure S8** Plot of apparent diffusion time extracted from the 2D1R fit of the 1 µs-started FCS curves against the peak threshold. The data were irregular and could not be described by equation (4). Dots represent experimental data. Data are shown of three independent experiments.

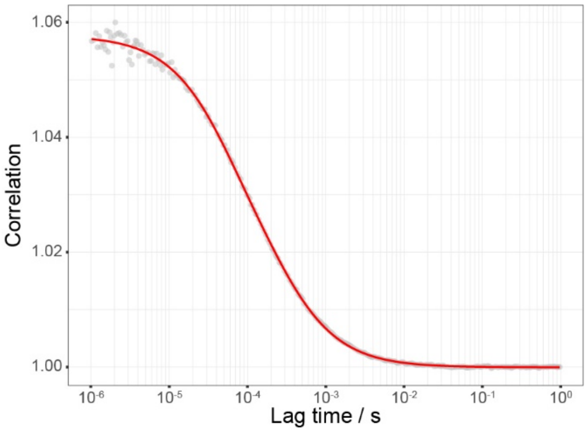


**Supplementary Figure S9** Example of the conventional FCS data and respective fit. The sample contains 5 nM Cy3B. Grey dots and red line represent experimental data and fitted curve.

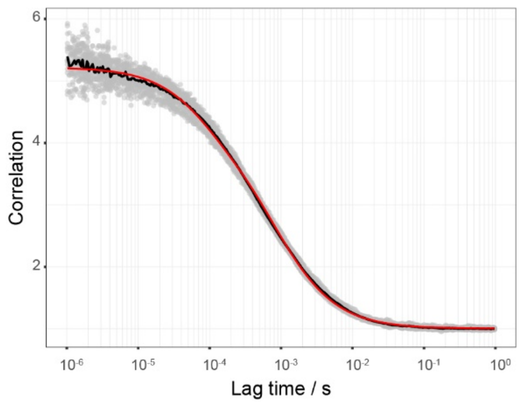


**Supplementary Figure S10** Example of the FCS data and respective fit for the Skp homo-trimerization. The sample contains 0.3 nM Skp D128C-Cy3B and 100 nM Skp. Grey dots, black line and red line represent experimental data, the data after averaging and fitted curve.

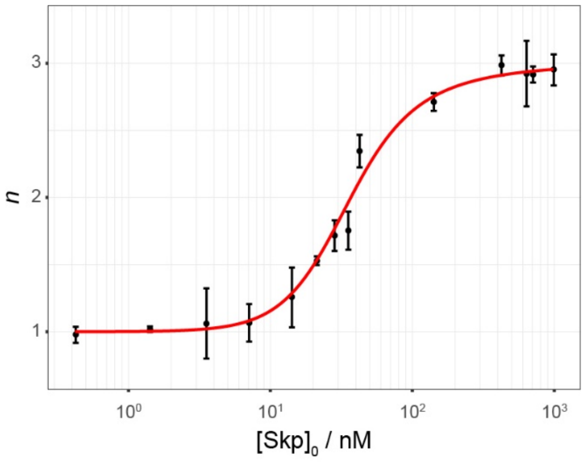


**Supplementary Figure S11** the titration curve of *n* as a function of $[Skp]_0$. Black dots and red line represent experimental data and fitted curves. Data are shown as mean±s.d. of three independent experiments.

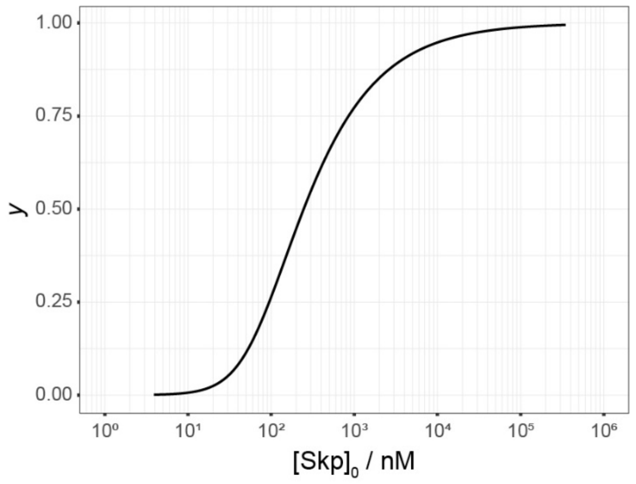


**Supplementary Figure S12** The titration curve of fraction of Skp in $Skp_3$ ($y$) as a function of $[Skp]_0$.

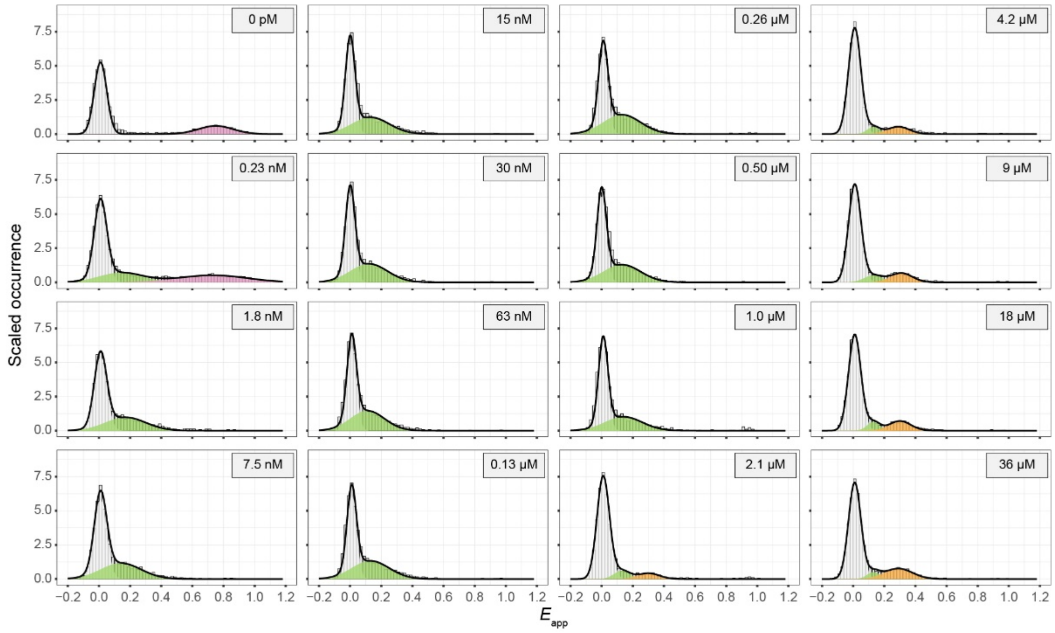


**Supplementary Figure S13** Intramolecular smFRET histogram of 50 pM OmpC G8C-D335C in Skp of different concentrations. Apo-OmpC, OmpC·$Skp_3$ and OmpC·$(Skp_3)_2$ are coloured in pink, green and amber. Zero-efficiency peak resulted from missing or inactivated acceptors is coloured in gray. All histograms were normalized and fitted by Gaussian distribution.

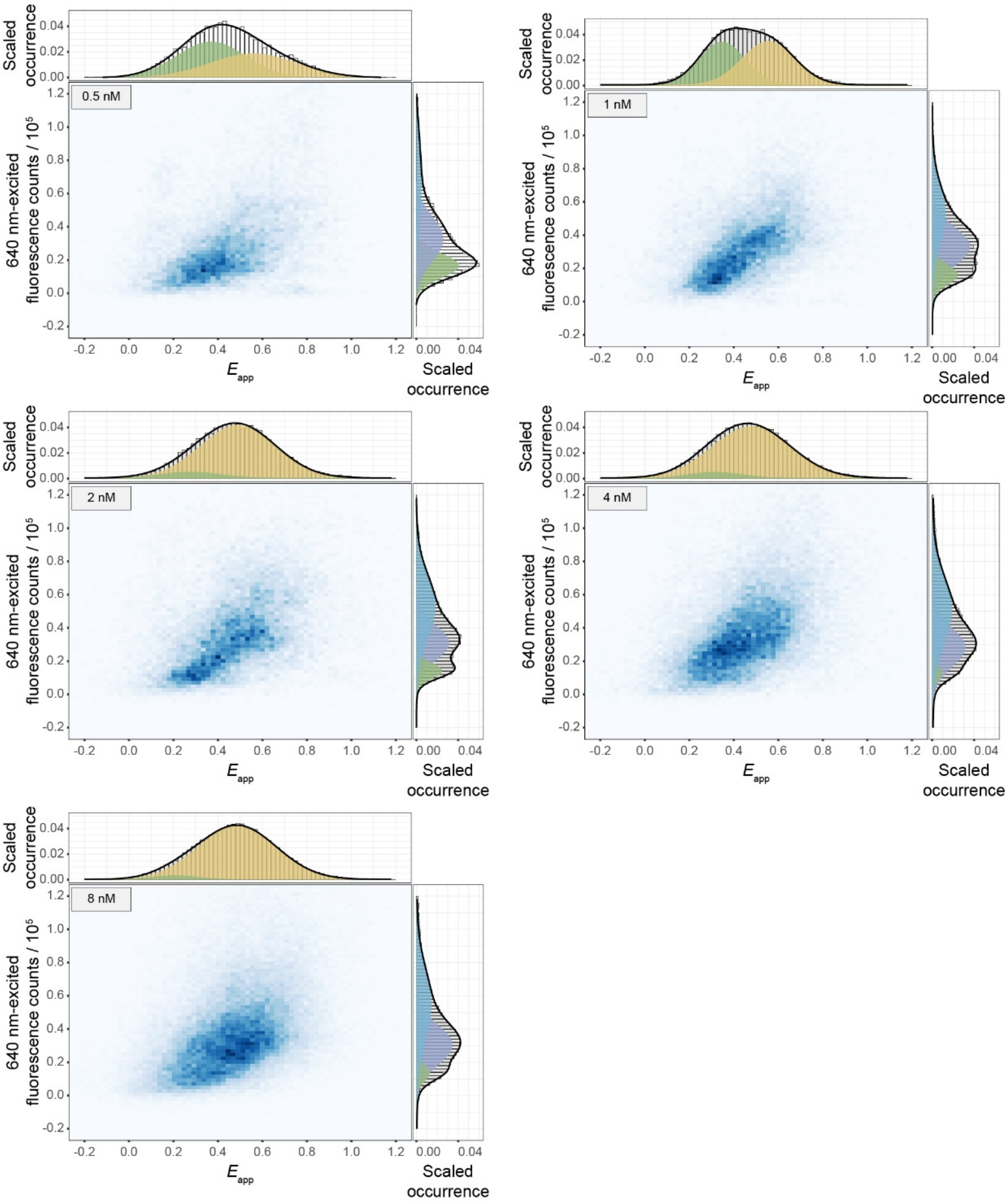


**Supplementary Figure S14** 2D histogram of immobilized OmpC G8C-AF555 in Skp D128C-AF647 of different concentrations. Histograms on the fluorescence counts dimension exhibited three peaks respectively positioned at $0.15\times10^5$ (green), $0.31\times10^5$ (indigo) and $0.56\times10^5$ (azure), which demonstrated that OmpC bound more than one Skp monomer. Histograms on the smFRET efficiency dimension could be fitted by two peaks respectively positioned at 0.30 (green) and 0.51 (amber), which suggests that the $E_{\mathrm{app}}$ was close when OmpC bound different numbers of fluorescent Skp monomers. All histograms were normalized and fitted by Gaussian distribution.

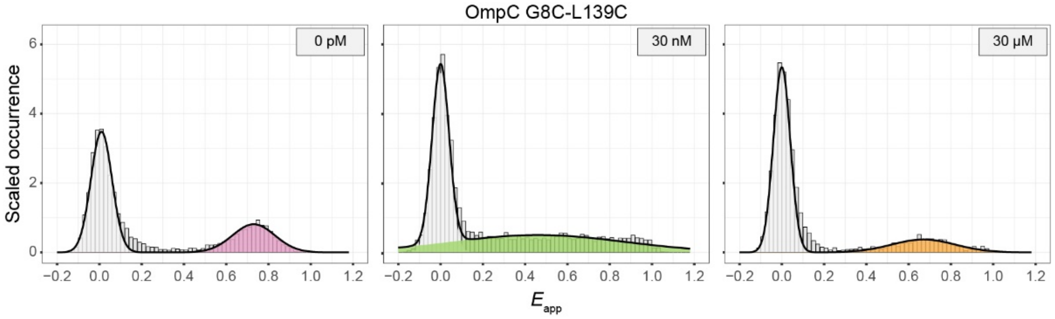


**Supplementary Figure S15** smFRET histogram of 50 pM OmpC G8C-L139C in 0 pM, 30 nM and 30 μM of Skp. Apo-OmpC, OmpC·$Skp_3$ and OmpC·$(Skp_3)_2$ are coloured in pink, green and orange. Zero-efficiency peak resulted from missing or inactivated acceptors is coloured in gray. All histograms were normalized and fitted by Gaussian distribution.

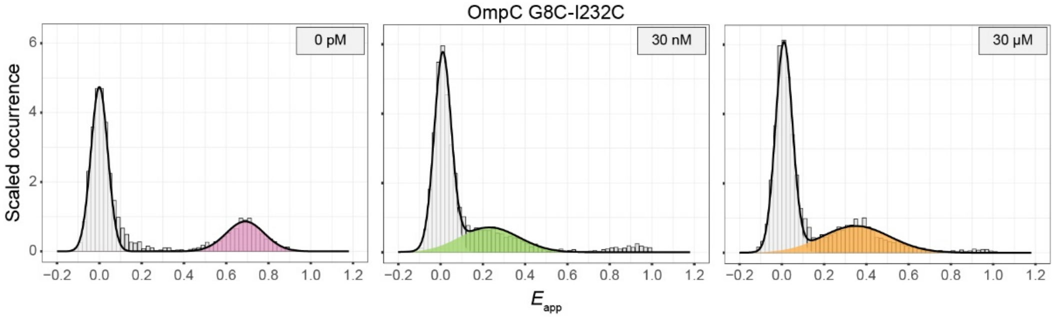


**Supplementary Figure S16** smFRET histogram of 50pM OmpC G8C-I232C in 0 pM, 30 nM and 30 μM of Skp. Apo-OmpC, OmpC·$Skp_3$ and OmpC·$(Skp_3)_2$ are coloured in pink, green and orange. Zero-efficiency peak resulted from missing or inactivated acceptors is coloured in gray. All histograms were normalized and fitted by Gaussian distribution.

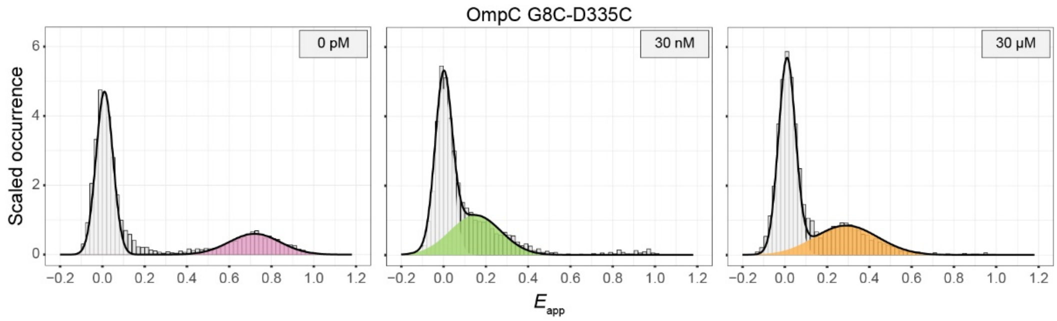


**Supplementary Figure S17** smFRET histogram of 50 pM OmpC G8C-D335C in 0 pM, 30 nM and 30 μM of Skp. Apo-OmpC, OmpC·$Skp_3$ and OmpC·$(Skp_3)_2$ are coloured in pink, green and orange. Zero-efficiency peak resulted from missing or inactivated acceptors is coloured in gray. All histograms were normalized and fitted by Gaussian distribution.

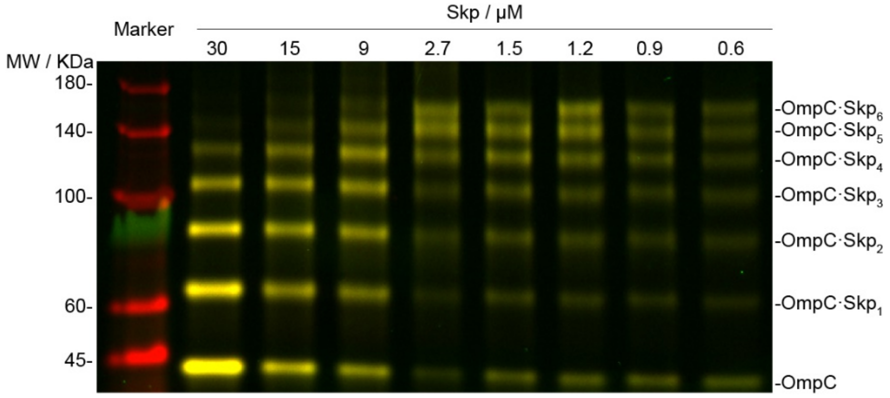


**Supplementary Figure S18** Cross-linked SDS-PAGE of Skp and intramolecular labelled OmpC G8C-D335C. The gel bands were assigned to the cross-linked OmpC·$Skp_n$ complexes according to their molecular weights.

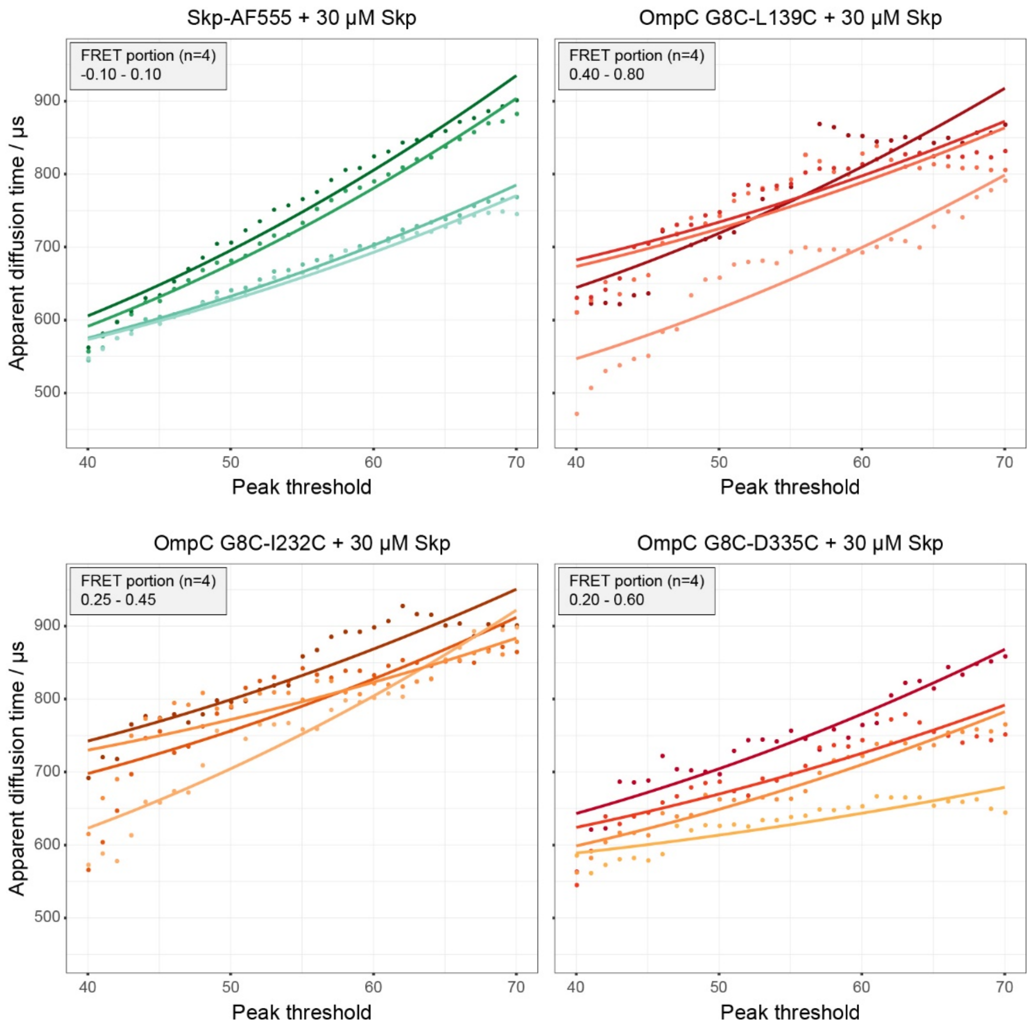


**Supplementary Figure S19** pscFCS of different dual-labelled OmpC mutants in 30 µM of Skp. Skp D128C-AF555 in 30 µM of Skp was used as reference. The calculated results are listed in Supplementary Table S2.

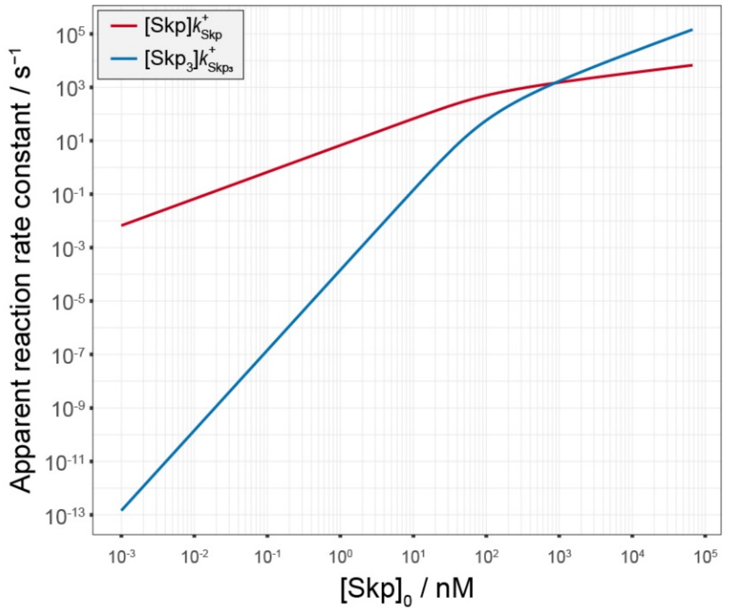


**Supplementary Figure S20** Estimation of reaction rate of Skp or $Skp_3$ to OmpC.

### Supplementary Tables

**Supplementary Table S1** Estimation of molecular weight and number of linked DSS molecules in intermolecular labelled OmpC and Skp in fluorescent SDS-PAGE

| Molecules | MW range (kD) | Theoretic MW (kD) | ΔMW (kD) | #DSS (Total) | #DSS (OmpC+Skp×n)[a] |
|---|---|---|---|---|---|
| OmpC·$Skp_6$ | 189 - 198 | 154 | 40 | 108 | 10+16×6 |
| OmpC·$Skp_5$ | 161 - 176 | 135 | 33 | 90 | 10+16×5 |
| OmpC·$Skp_4$ | 134 - 141 | 116 | 22 | 59 | 10+12×4 |
| OmpC·$Skp_3$ | 112 - 121 | 97 | 19 | 53 | 10+14×3 |
| OmpC·$Skp_2$ | 90 - 93 | 78 | 14 | 37 | 10+13×2 |
| OmpC·$Skp_1$ | 65 - 71 | 59 | 10 | 27 | 10+17×1 |
| $Skp_3$ | 65 - 71 | 57 | 10 | 27 | 0+9×3 |
| OmpC | 43 - 44 | 40 | 3.5 | 10 | 10+0 |
| $Skp_2$ | 41 - 42 | 38 | 3.6 | 10 | 0+5×2 |
| Skp | 16 - 20 | 19 | -0.9 | -3 | 0+0 |

a  Proposed possible numbers of DSS molecules among protein molecules

**Supplementary Table S2** hydrodynamic radius of different molecules derived from pscFCS

| **Subpopulation** | **Labelling sites** | **Diffusion time / µs** | **Radius / nm** |
|---|---|---|---|
| $Skp_3$ | Skp D128C | 459±19 | 3.3[a]±0.1 |
| $OmpC{\cdot}Skp_3$ | OmpC G8C-D335C | 549±27 | 3.9±0.2 |
| $OmpC{\cdot}(Skp_3)_2$ | OmpC G8C-L139C | 527±77 | 3.8±0.6 |
| $OmpC{\cdot}(Skp_3)_2$ | OmpC G8C-I232C | 592±81 | 4.3±0.6 |
| $OmpC{\cdot}(Skp_3)_2$ | OmpC G8C-D335C | 533±16 | 3.8±0.1 |

a value from reference 34 in the main text.
